# A new class of natural anthelmintics targeting lipid metabolism

**DOI:** 10.1101/2024.12.09.627452

**Authors:** Hala Zahreddine Fahs, Fathima S. Refai, Suma Gopinadhan, Yasmine Moussa, Hin Hark Gan, Yamanappa Hunashal, Gennaro Battaglia, Patricia G. Cipriani, Claire Ciancia, Nabil Rahiman, Stephan Kremb, Xin Xie, Yanthe E. Pearson, Glenn L. Butterfoss, Rick M. Maizels, Gennaro Esposito, Antony P. Page, Kristin C. Gunsalus, Fabio Piano

## Abstract

Parasitic helminths are a major global health threat, infecting nearly one-fifth of the human population and causing significant losses in livestock and crops. Resistance to the few anthelmintic drugs is increasing. Here, we report a set of avocado fatty alcohols/acetates (AFAs) that exhibit nematocidal activity against four veterinary parasitic nematode species: *Brugia pahangi*, *Teladorsagia circumcincta* and *Heligmosomoides polygyrus*, as well as a multidrug resistant strain (UGA) of *Haemonchus contortus.* AFA shows significant efficacy in *H. polygyrus* infected mice. In *C. elegans*, AFA exposure affects all developmental stages, causing paralysis, impaired mitochondrial respiration, increased reactive oxygen species production and mitochondrial damage. In embryos, AFAs penetrate the eggshell and induce rapid developmental arrest. Genetic and biochemical tests reveal that AFAs inhibit POD-2, encoding an acetyl CoA carboxylase, the rate-limiting enzyme in lipid biosynthesis. These results uncover a new anthelmintic class affecting lipid metabolism.

## Introduction

More than 1.5 billion people are infected with helminth parasites, primarily affecting marginalized populations in low- and middle-income countries, according to the World Health Organization^1^. These parasites are transmitted by skin-penetrating larvae or by eggs that are shed in human faeces and contaminate soil in areas where sanitation is poor^2^. Helminth infections also have a devastating impact on companion animals, livestock production globally and are damaging to all major crops, causing huge economic losses estimated at over $80 billion per year^3–6^.

Helminths compete with insects as the most evolutionarily successful animal clade on the planet, having colonized almost all known environments, including several extreme ones^7,8^. Parasitic helminths use animals or plants as their host, resulting in both direct and indirect negative impacts on public health, agricultural systems, and the ecology and conservation of wild species^7,9,10^. Helminth infections are particularly challenging to treat without causing toxicity to mammalian hosts, due to strong similarities in their physiology and a multitude of mechanisms to block uptake, metabolize or expel exogenous poisons.

Despite the global challenges posed by animal and human helminth infections, the number of drugs used to treat them is limited to only a few chemical classes and target pathways. These include the benzimidazoles, targeting beta tubulin (e.g. albendazole); the macrocyclic lactones, targeting glutamate-gated chloride channels (e.g. ivermectin); the imidazothiazoles, targeting nicotinic acetylcholine receptors (e.g. levamisole); the pyrazinoisoquinolones, targeting calcium channels (e.g. praziquantel); the salicylanilides, uncouplers of oxidative phosphorylation (e.g. closantel); and piperazine, a GABA receptor agonist^11^. As drug resistance among parasitic helminths is on the rise^11–14^, new chemical classes with novel mechanisms of action are thus urgently needed to overcome the threat of resistance^11^. Nature has been a valuable source of anthelmintic compounds, exemplified by the discovery in the 1970s of avermectin, a natural product of the soil bacterium *Streptomyces avermitilis*, and its widely used derivative ivermectin^15^. In particular, plants have evolved a multiplicity of defense molecules against pathogens and therefore are a likely source of new, natural anti-nematode agents.

Direct high-throughput screening (HTS) of drugs for activity against parasitic helminths is challenging because their life cycle requires a developmental phase in the host, adding layers of complexity to the screening process^16^. The free-living nematode *Caenorhabditis elegans* has already been established as a model for anthelmintic drug discovery due to its ease of laboratory culture, amenability to HTS for rapid whole-organism phenotypic screening, and the availability of extensive forward and reverse genetic methods^16–18^. Molecular genetic approaches, combined with chemical perturbation, provide a powerful toolbox for *in vivo* mode of action studies to identify molecular targets^18–20^. Moreover, parallel HTS across multiple species including *Pristionchus pacificus* could help identify candidates with broad-spectrum activity. Counter-screening for low toxicity in human cells can then help focus further efforts on potential lead compounds that are readily tolerated by humans.

Here, we describe a multispecies HTS approach using *C. elegans* and *P. pacificus* to screen a commercial small molecule library containing FDA-approved drugs and natural products for potential broad-spectrum anthelmintic activity (Supplementary data 1). Most molecules in this library are expected to be well tolerated in humans, which we confirmed by measuring toxicity in a human cell line. Among the bioactive compounds identified in the screen was a group of structurally similar 17-carbon fatty alcohol compounds present in extracts of the avocado *Persea americana*. We found that these compounds, which we collectively refer to as avocado-derived fatty alcohols/acetates (AFAs), elicit dose-dependent lethality in parasitic nematodes and target *C. elegans* POD-2, an acetyl-CoA carboxylase (ACC) that is the rate limiting enzyme in lipid biosynthesis.

## Results

### Novel class of anthelmintics

To identify new compounds with potential as anthelmintics, we screened a library of 2,300 small molecules containing FDA-approved drugs, other known bioactive compounds, and natural products (Spectrum Collection, MicroSource Discovery Systems, Inc.) (Supplementary data 1). Since parasitic nematodes are not amenable to high throughput screening (HTS) due to their complex life cycle, to identify potential broad-spectrum anthelmintics we looked for compounds that affect both of two distantly related free-living model species, *C. elegans* and *P. pacificus* (Fig. 1a). To avoid non-specific cytotoxicity, especially in human cells, we additionally required low toxicity in U2-OS human bone osteosarcoma cells that we and others use routinely for image analysis due to their flattened morphology (Supplementary Fig. 1)^21,22^.

**Fig. 1.**
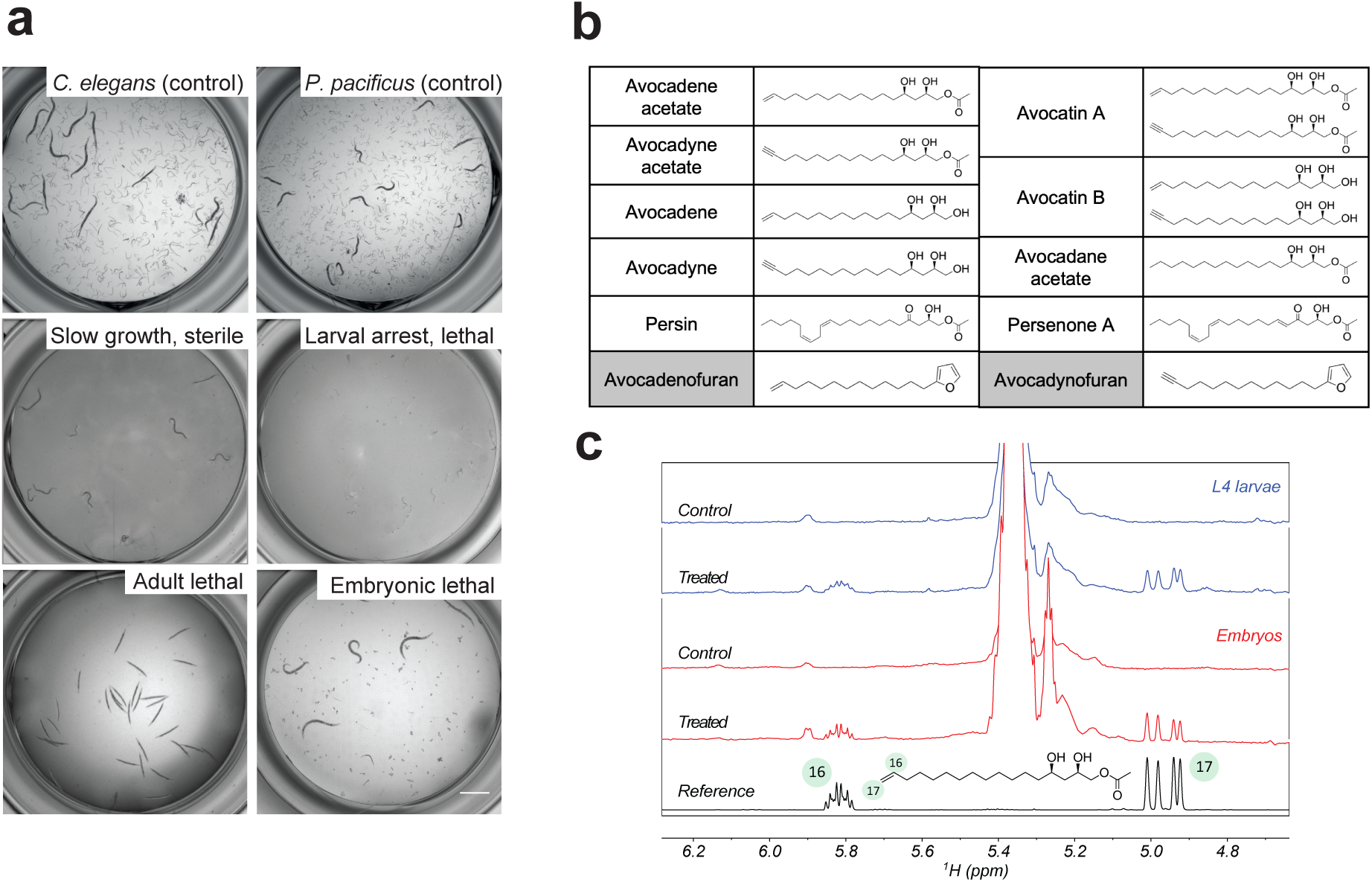
A screen using *C. elegans* and *P. pacificus* identifies novel anthelmintic compounds. **a,** Treatment with AFA hits induced slow growth, sterility, larval lethality, larval arrest, adult lethality or paralysis or embryonic lethality (*n* = 3 biological replicates (BRs) with 2 technical replicates (TRs) each) (scale bar = 1 mm). **b,** Chemical structure of avocado-derived compounds. All except the furans show anthelmintic activity against *C. elegans* and *P. pacificus*. **c,** Incorporation of avocadene acetate in *C. elegans* embryos and larvae as detected by the ^1^H NMR spectrum of the pure reference compound compared with the corresponding spectra of CHCl_3_ fraction extracts from embryos and L4 larvae, with and without treatment with 20 μM or 10 μM avocadene acetate, respectively.

The primary screen was performed by treating synchronized L1 larvae with 10 µM compounds and incubating for 5-7 days (see methods, Supplementary data 2) and resulted in the identification of a family of related compounds isolated from the avocado *Persea americana* that cause a variety of severe phenotypes in *C. elegans* (Fig. 1a). These AFA compounds represent structurally similar 17-carbon fatty alcohols and their acetate variants (Fig. 1b): *avocadene* ((2R,4R)-1,2,4-trihydroxyheptadec-16-ene), *avocadyne* ((2R,4R)-1,2,4-trihydroxyheptadec-16-yne), *avocadene acetate* ((2R,4R)-2,4-dihydroxyheptadec-16-en-1-yl acetate), *avocadyne acetate* ((2R,4R)-2,4-dihydroxyheptadec-16-yn-1-yl acetate), *avocatin B* (a mixture of avocadene and avocadyne), and *avocatin A* (a mixture of their respective acetate forms).

Nematodes are known to be highly resistant to harsh environmental conditions. Infective and adult stages are surrounded by a thick protective cuticle and embryos are protected by multiple layers that include a chitin layer, a vitelline layer and a chondroitin proteoglycan layer^23^. Using the model *C. elegans*, we asked if AFAs accumulate inside embryos or post-embryonic stages. Analysis of embryo and larval extracts by proton NMR revealed that both embryos and larvae contained significant amounts of AFAs following 10-20 µM treatments, as shown by the presence of the characteristic signals of the terminal olefinic group of avocadene acetate (Fig. 1c).

To characterize the activity profile of AFAs, we assayed them at different concentrations (from 1 µM to 100 µM) in liquid culture across *C. elegans* developmental stages, in 3 biological replicates (*n* = 3) and 4 technical replicates (*n* = 4) (Fig. 2). We found that all six of these (avocadene, avocadyne, their acetate forms, and the two mixtures avocatin A and B) yielded a concentration-dependent toxic effect on L1 larval development (Fig. 2a), adult survival (Fig. 2b) and egg hatching (Fig. 2c, Supplementary movie 1, Supplementary movie 2). In contrast, two structurally related compounds, avocadenofuran and avocadynofuran, which contain a 17-carbon chain but differ in having a furan ring (Fig. 1b), showed no bioactivity in any of our assays at concentrations up to 100 µM (Fig. 2a-d). Comparison of LD_50_ values showed that avocadene acetate, avocadyne acetate, and avocatin A exhibited similar potency and showed stronger effects than the non-acetate forms (Fig. 2d).

**Fig. 2.**
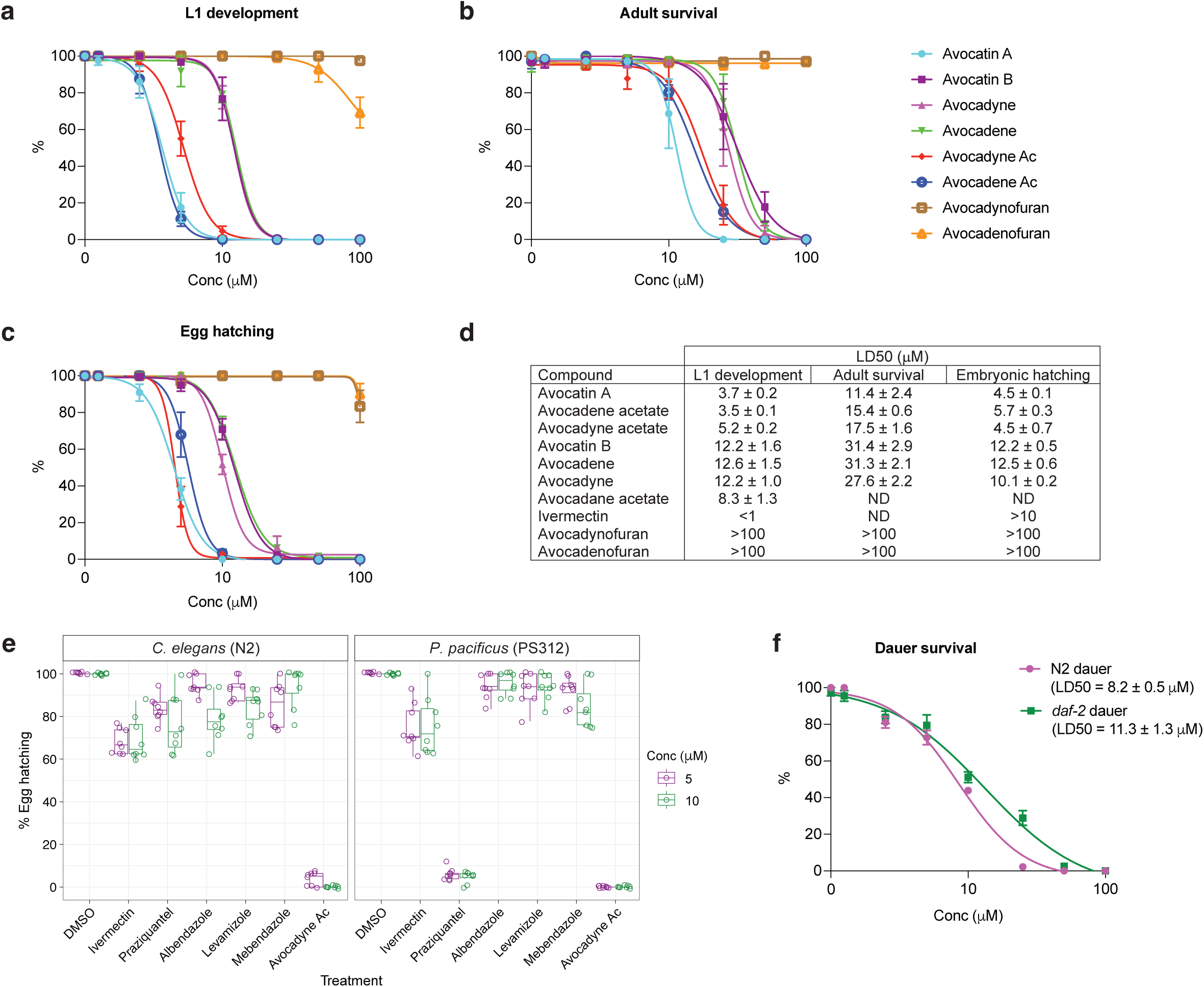
AFAs impair development and survival in *C. elegans* and *P. pacificus*. **a-c,** Dose-response curves of the 8 AFA compounds. **a,** Percentage of animals developing past the L1 stage after 5 days of treatment beginning at the L1 stage. **b,** Percentage of adults surviving after 48 hours of treatment. **c,** Percentage of eggs hatched after 48 hours of treatment. **d,** Calculated LD_50_ values (concentration at which development or survival was reduced by 50%) of AFA compounds and ivermectin in each assay. **e,** Percentage of eggs hatched in N2 and PS312 after 48 hours of treatment with existing anthelmintics and avocadyne acetate at 5 and 10 µM. Lower and upper bounds of the box correspond to the first and third quartiles (25th and 75th percentile), whiskers extend to the largest or smallest value no further than 1.5 times the inter-quartile range (IQR). **f,** Dose-response curves of avocadyne acetate on the percentage survival of N2 dauers and *daf-2* dauers upon 48 hours of exposure to the compound. Each data point represents mean ± SEM; for (**a-c,f**) *n* = 3 BRs with 4 TRs each with >50 worms/concentration; for (**e**) *n* = 3 BRs with 3 TRs each with >100 individual eggs/concentration. In all experiments, DMSO was used as a negative control. Source data are provided as a Source Data file.

To further explore this class of compounds, we asked if related molecules would have similar effects. We found that two other purified molecules extracted from Avocado, persenone A ((2R,5E,12Z,15Z)-2-hydroxy-4-oxoheneicosa-5,12,15-trienyl acetate) and persin ((2R, 12Z,15Z)-2-hydroxy-4-oxoheneicosa-12,15-dienyl acetate), as well as a fully saturated synthetic molecule, avocadane acetate ((2R,4R)-2,4-dihydroxyheptadecan-1-yl acetate) (Fig. 1b), had similar activities (Supplementary Fig. 2). Since persin and its derivatives are known to be toxic to animals^24^, we focused on the novel AFAs we found in the screen.

The egg hatching assay was performed on harvested embryos, and we confirmed the presence of AFAs within treated embryos using NMR (Fig. 1c), demonstrating that these lipid alcohols and acetates can cross the eggshell barrier. To compare potency, we measured *C. elegans* and *P. pacificus* egg hatching in the presence of avocadyne acetate or a selection of other anthelmintics (Fig. 2e). We found that albendazole, mebendazole, and levamisole had no effect on egg hatching in either species; praziquantel (used to treat infections of the skin-penetrating worm *Schistosoma*) reduced egg hatching by 25% (*n* = 517; unpaired t-test, *p* < 0.0001) in *C. elegans* and by 90% (*n* = 416; unpaired t-test, *p* < 0.0001) in *P. pacificus* at both 5 µM and 10 µM, and ivermectin decreased egg hatching by only 25% in both species (*n* = 594 and 751, respectively; unpaired t-test, *p* < 0.0001). In contrast, avocadyne acetate led to 100% inhibition of egg hatching in both worm models at 10 µM (*n* = 466 in *C. elegans* and *n* = 447 in *P. pacificus*). AFAs thus show a rare ability among known anthelmintics to directly interfere with embryogenesis (independently of the maternal provisioning of fat molecules during oogenesis), which would be highly favorable for the treatment of parasitic nematodes.

Nematodes such as *C. elegans* can enter a dauer stage, during which they reduce their metabolism in order to adapt to harsh environments such as dryness, reduced nutrients, and altered temperature. The dauer stage is thought to resemble the infective L3 stages of many parasitic nematodes^25^. To determine whether dauer larvae are also sensitive to AFAs, we treated dauers from both wild-type N2 (obtained by starvation) and the *daf-2(m41)* strain (which induces dauers at 25°C)^19^ with avocadyne acetate for 48 h at 25°C, and found that it elicited lethality in N2 and *daf-2(m41)* dauer larvae with an LD_50_ of 8.5 µM and 13.5 µM, respectively (Fig. 2f).

### AFAs show a new mode of action

To test the effect of the AFA compounds on parasitic nematodes, we conducted assays in four veterinary parasites: *H. contortus*, *T. circumcincta*, *H. polygyrus* and *B. pahangi*. Avocatin B at both 10 µM and 50 µM induced strong larval lethality in all three parasitic species tested: *H. contortus*, *T. circumcincta*, and *H. polygyrus* (Table 1, Supplementary table 1). Furthermore, AFAs were also active in the multidrug resistant *H. contortus* UGA larvae and eggs at 10 µM and 50 µM, respectively (Table 1). Importantly, avocatin A not only is effective against L3 and microfilarial stages, but also acts as macrofilaricidal, as it kills adult *B. pahangi* starting at 8-10 µM, with adult females being more sensitive than males (Table 2). In *H. contortus*, while all six bioactive AFA compounds from the primary screen impaired egg hatching and caused larval lethality, the acetate forms elicited the strongest phenotypes (Supplementary table 1). Together with results from the primary screen, this analysis confirmed that the acetate forms induce the highest potency effect. Therefore, we performed the subsequent experiments using either avocadene acetate or avocatin A, which is a mix of the two acetate forms.

**Table 1.** AFAs are effective against a multi-drug resistant parasitic nematode strain. Rates of larval lethality of parasitic worms after treatment with avocatin A or B. *T. circumcincta* L2 counts are derived from biological triplicates (*n* = 3) and *H. contortus (UGA)* L2 counts from biological duplicates (*n* = 2). *H. polygyrus* L2 and *H. contortus (UGA)* eggs counts are derived from a singlet observation.

|  | Conc. | *T. circumcincta* L2 |  | *H. polygyrus* L2 |  | *H. contortus* (UGA) L2 |  | *H. contortus* (UGA) eggs |  |
| --- | --- | --- | --- | --- | --- | --- | --- | --- | --- |
|  |  | Total | % lethality | Total | % lethality | Total | % lethality | Total | % lethality |
| DMSO | 1% | 32 | 18.8 | 52 | 21.2 | 215 | 30.1 ± 2.3 | 113 | 33.6 |
| Avo B | 10µM | 119 | 87.6 ± 5.1 | 40 | 100 | 223 | 79.4 ± 8 | 89 | 31.5 |
|  | 50µM | 119 | 100 ± 0 | 69 | 100 | 209 | 100 ± 0 | 30 | 100 |
| Avo A | 10µM | – | – | 55 | 100 | 223 | 97.1 ± 2.9 | 66 | 36.4 |
|  | 50µM | – | – | 76 | 100 | 162 | 100 ± 0 | 75 | 100 |

**Table 2.** AFAs are effective against filarial nematode *B. pahangi*. Rates of larval lethality of *B. pahangi* adults, females and males, L3 and microfilariae (mfs) after treatment with avocatin A. *B. pahangi* adult counts are derived from 5 biological replicates (*n* = 5). Females, males, L3 and microfilaria are from a singlet observation.

| *B. pahangi* | DMSO | 2 µM | 4 µM | 8 µM | 10 µM | 15 µM | 20 µM |
| --- | --- | --- | --- | --- | --- | --- | --- |
| Total adults | 40 | 32 | 32 | 31 | 36 | 34 | 37 |
| % lethality/paralysis | 0 | 0 | 0 | 2.5 ± 2.2 | 15 ± 6.1 | 46.3 ± 13.6 | 100 ± 0 |
| Total females | 10 | 10 | 10 | 10 | 10 | 10 | 10 |
| % lethality/paralysis | 0 | 0 | 0 | 0 | 20 | 50 | 100 |
| Total males | 10 | 10 | 10 | 10 | 10 | 10 | 10 |
| % lethality/paralysis | 0 | 0 | 0 | 20 | 30 | 90 | 100 |
| Total L3 | 61 | – | – | – | 68 | – | 69 |
| % lethality | 0 | – | – | – | 100 | – | 100 |
| Total Mfs | 104 | 154 | 131 | 150 | 126 | 147 | 163 |
| % lethality/paralysis | 1.9 | 5.2 | 9.2 | 100 | 100 | 100 | 100 |

To test the possibility that AFAs represent a new class of anthelmintics, we screened them in *C. elegans* mutants that are resistant to the known anthelmintics ivermectin/abamectin, levamisole, benzimidazoles and amino-acetonitrile derivative (AAD) monepantel. All mutant strains were susceptible to AFA toxicity at the same level as wild type N2 when treated at the L1 stage, indicating that AFA compounds act through a different mechanism (Table 3, Supplementary Fig. 3). To further characterize the mode of action of AFAs, we performed random mutagenesis in *C. elegans* using ethylmethanesulfonate (EMS). Despite performing several large rounds of screening and testing the resistance of ∼10 million genomes, we were unable to recover any strain with a reproducible level of resistance. The results of this forward genetic screen suggest either that evolving resistance to AFAs may require changes in multiple loci, or that their anthelmintic effects arise from physical damage to cellular structures.

**Table 3.**
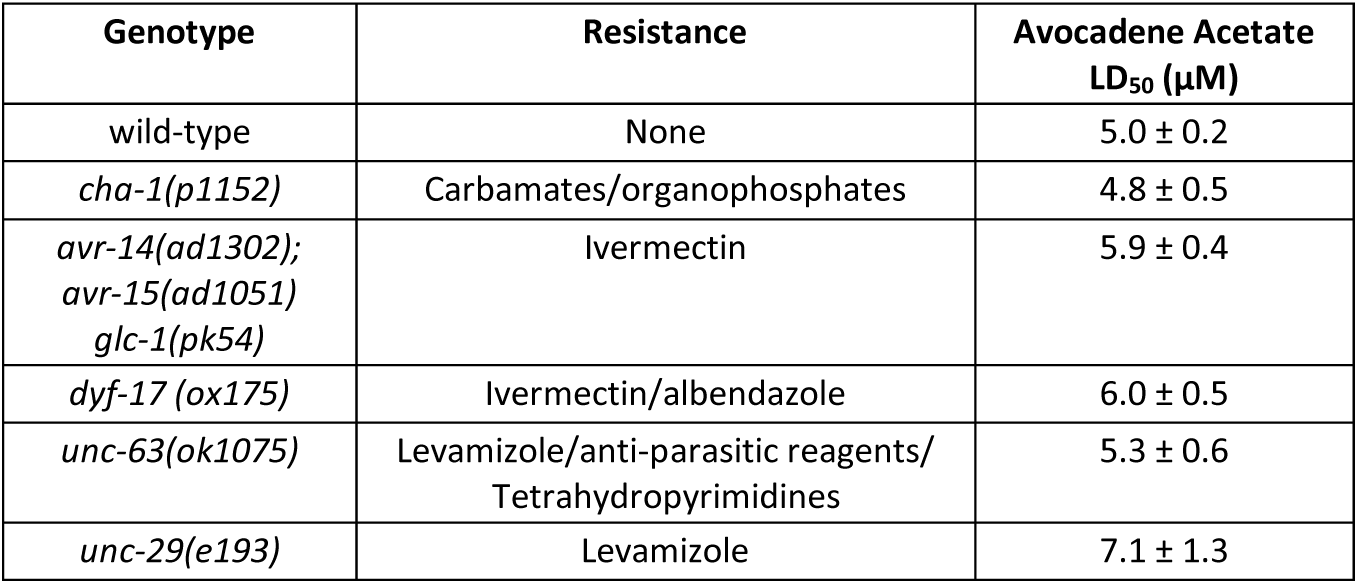
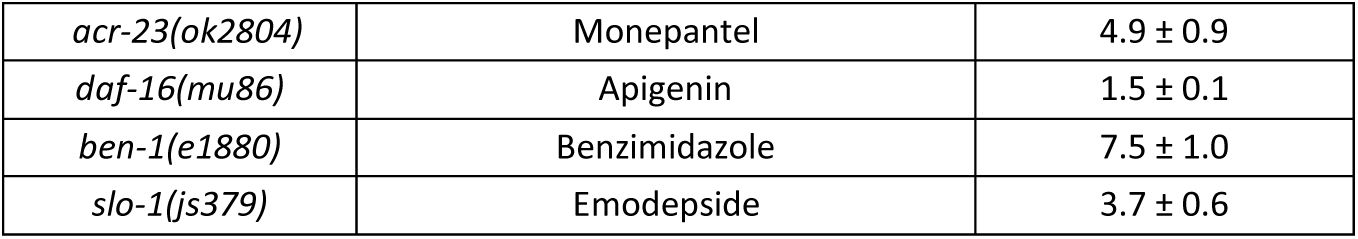
AFAs. Calculated LD_50_ values of AFA compounds in the anthelmintic-resistant *C. elegans* mutant strains treated at L1 stage. Data derived from 3 biological replicates (*n* = 3).

### AFAs are active *in vivo*

To determine whether the anthelmintic effects of AFAs tested directly on nematodes would translate to an *in vivo* parasitic nematode system, we used mice infected by *H. polygyrus,* a model for gastrointestinal infection^26^. We tested the effect of Avocatin A on infected mice that were pre-inoculated and allowed to develop a high titer of infection for a week. Infected mice treated with Avocatin A showed consistent and significant reduction compared with a placebo control in both *H. polygyrus* fecal egg count (FEC) and in the number and health of adult worms dissected from the mice on day 21 (*p* < 0.01 and *p* < 0.05, unpaired t-test) (Fig. 3). Three of the ten tested mice were essentially cured of the *H. polygyrus* infection within the two-week trial period and four out of ten produced no eggs.

**Fig. 3.**
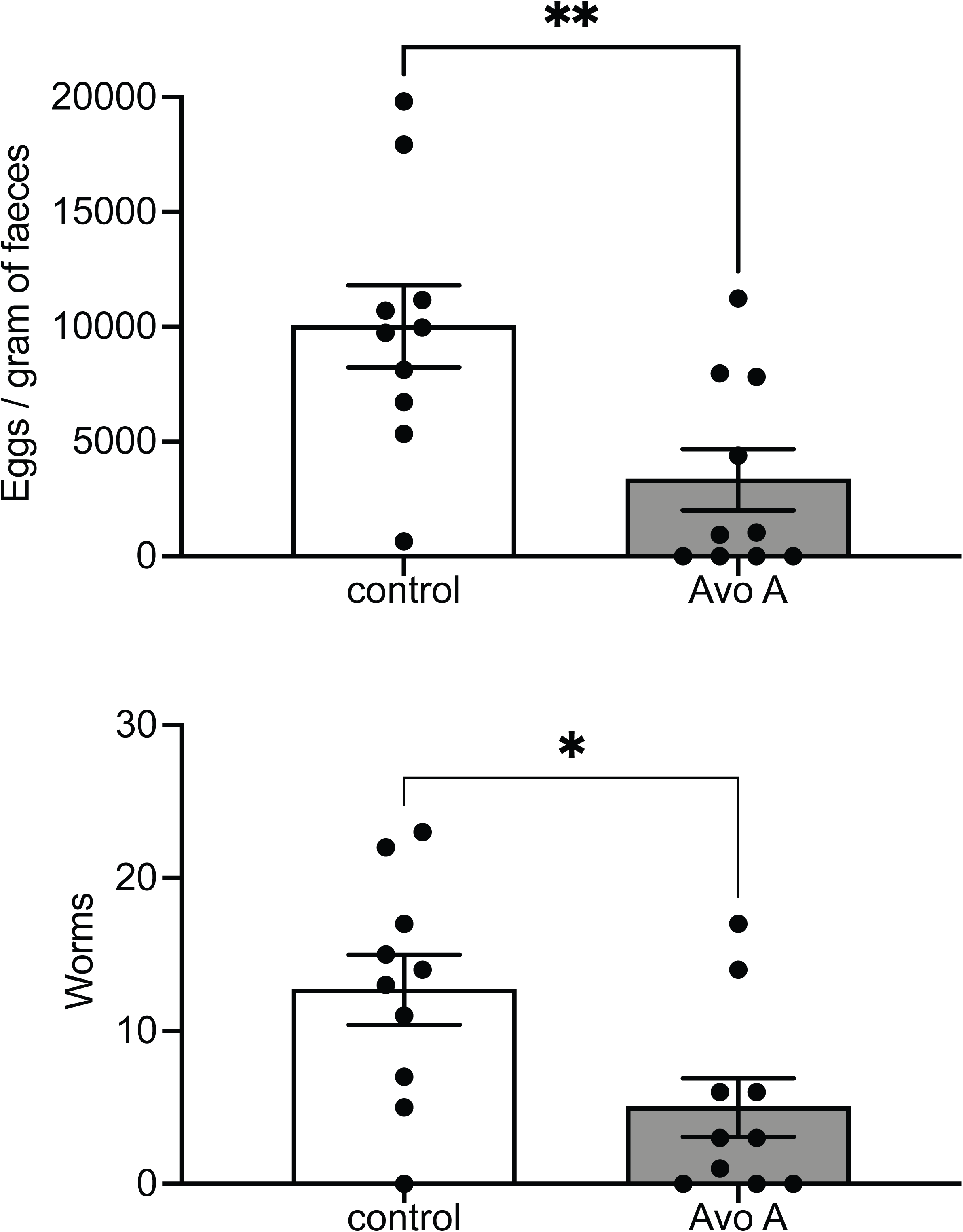
AFAs are active *in vivo.* Eggs per gram of feces and adult worm counts are reduced after avocatin A (100mg/kg) treatment in mice infected with *H. polygyrus.* C57BL/6 mice were infected with 200 *H. polygyrus* larvae and were given Avocatin A or control vehicle on alternate days from day 9 to 19; egg counts and adult worm numbers were enumerated at day 21 post infection. Data represent pooled results from two independent experiments, each with 5 mice per group, and analyzed by unpaired two-tailed t-test (*p* = 0.0076 for eggs and *p* = 0.0188 for worms). Where applicable, data represents mean ± SEM. Cells with ‘–’ indicate that the assay was not performed. Source data are provided as a Source Data file.

### AFAs inhibit respiration in *C. elegans*

To explore the mode of action of AFAs, we examined the phenotypic effects of treating *C. elegans* embryos and adults with avocadene acetate, one of the most potent AFA compounds. Adults displayed dose-dependent paralysis and inhibition of pharyngeal pumping within ∼5 minutes of treatment (Fig. 4a-c, Supplementary Fig. 4). In embryos, acute treatment with 50 µM led to developmental arrest of all embryonic stages within ∼30 minutes (Fig. 5f, Supplementary Fig. 5). To further characterize the paralysis, we measured oxygen consumption rates (OCR) in whole nematodes at the L4 stage, using a Seahorse XFe96 Analyzer (Fig. 4f)^27^. Basal OCR, a measure of baseline mitochondrial respiration, was significantly lower in worms treated with sub-lethal concentrations of avocadene acetate in comparison with DMSO controls (unpaired t-test, *p* < 0.0001) (Fig. 4f). We next measured maximal OCR in the presence of carbonyl cyanide-p-trifluoromethoxy phenylhydrazone (FCCP). FCCP is an uncoupler of mitochondrial oxidative phosphorylation that collapses the mitochondrial inner membrane proton gradient, which inhibits ATP synthesis and causes the electron transport chain to work at full capacity^27^. These experiments showed that worms treated with sub-lethal concentrations of avocadene acetate exhibited significantly lower maximal respiration and spare capacity (maximal OCR - basal OCR) in comparison with controls (unpaired t-test, *p* < 0.001 and *p* < 0.0001) (Fig. 4f, Supplementary Fig. 6a).

**Fig. 4.**
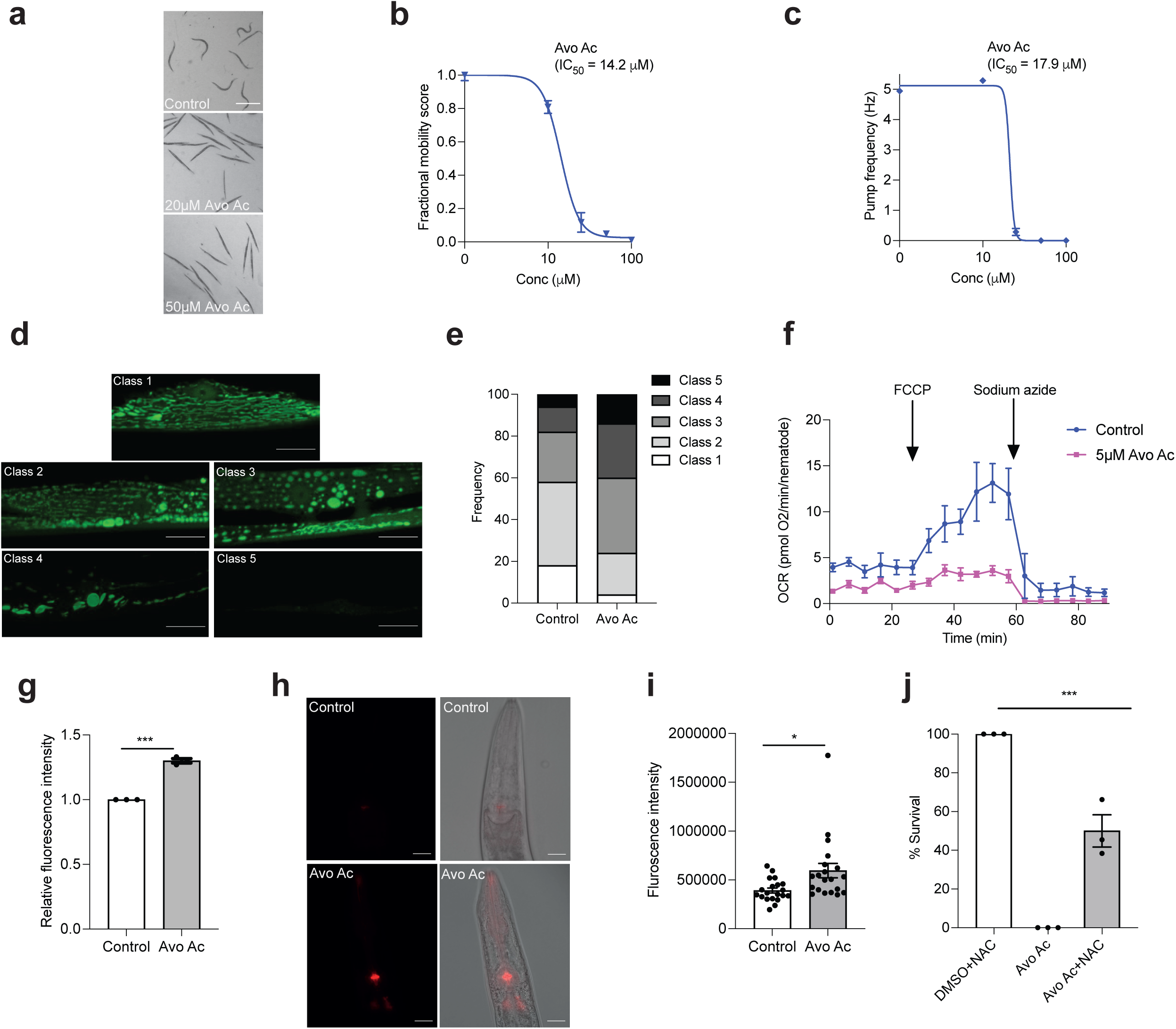
Avocadene acetate causes paralysis and ROS production in *C. elegans*. **a,** Stereoscope live imaging demonstrating the paralysis phenotype upon avocadene acetate treatment (*n* = 3 BRs) (scale bar = 1 mm). **b,** Dose response curve of motility of young adult worms treated with varying concentrations of avocadene acetate for 5 minutes. **c,** Dose response curve of pharyngeal pump frequency of young adult worms treated with avocadene acetate. Datapoints in (**b and c**) represent mean ± SEM; *n* = 3 BRs with 5 TRs each for **b** and *n* = 3 BRs with > 20 individual worms per concentration for **c**. **d,** Representative images of body wall muscle mitochondria in *C. elegans s*train SJ4103, illustrating the five morphological classes of progressive mitochondrial deterioration: Class 1, tubular network (similar to wild type); Classes 2- 4, altered morphology with blebbing, fragmentation, and enlarged, rounded appearance (Class 2, abundant mitochondria; Class 3, fewer mitochondria; Class 4, very few mitochondria); Class 5, extremely few or no mitochondria detected (scale bar = 10 µm). L1 animals were treated for 48-hours with 0.5% DMSO (control) or 5 µM avocadyne acetate. **e,** Distribution of mitochondrial defects in control and treated animals, *n* = 12). Around 50 muscles from each group from two independent experiments were analyzed (*p* = 0.005, two sided Chi-square test with df = 4). Controls for all treatments were DMSO. **f,** Kinetic oxygen consumption rate (OCR) measuring basal respiration, FCCP-driven OCR (maximal respiration) and sodium azide-driven OCR (non-mitochondrial respiration). Each time point represents the average OCR±SEM of five individual wells, expressed as pmol/min/animal. **g,** Intracellular ROS in 10 µM avocadene acetate treated animals (L1s treated for 48 hours) normalized to controls, using CM-H2DCFDA method. Three independent experiments were performed (unpaired two-tailed t-test, *p* < 0.0001). **h,** Representative images of Mitosox fluorescence in the posterior pharyngeal region (left) with DIC overlay (right) in control and 10 µM avocadene acetate treated worms (L1s for 42 hours) (scale bar = 20 µm). **i,** Mitosox fluorescence quantification using ImageJ software (*n* = 20; unpaired two-tailed t-test, *p* = 0.01). **j,** Pre-treatment with 10 mM N-acetyl cysteine (NAC) partially rescued lethality induced by 15 µM avocadene acetate on L4 stage animals (*n* = 3 BRs, one-way ANOVA, *p* < 0.0001). Source data are provided as a Source Data file.

**Fig. 5.**
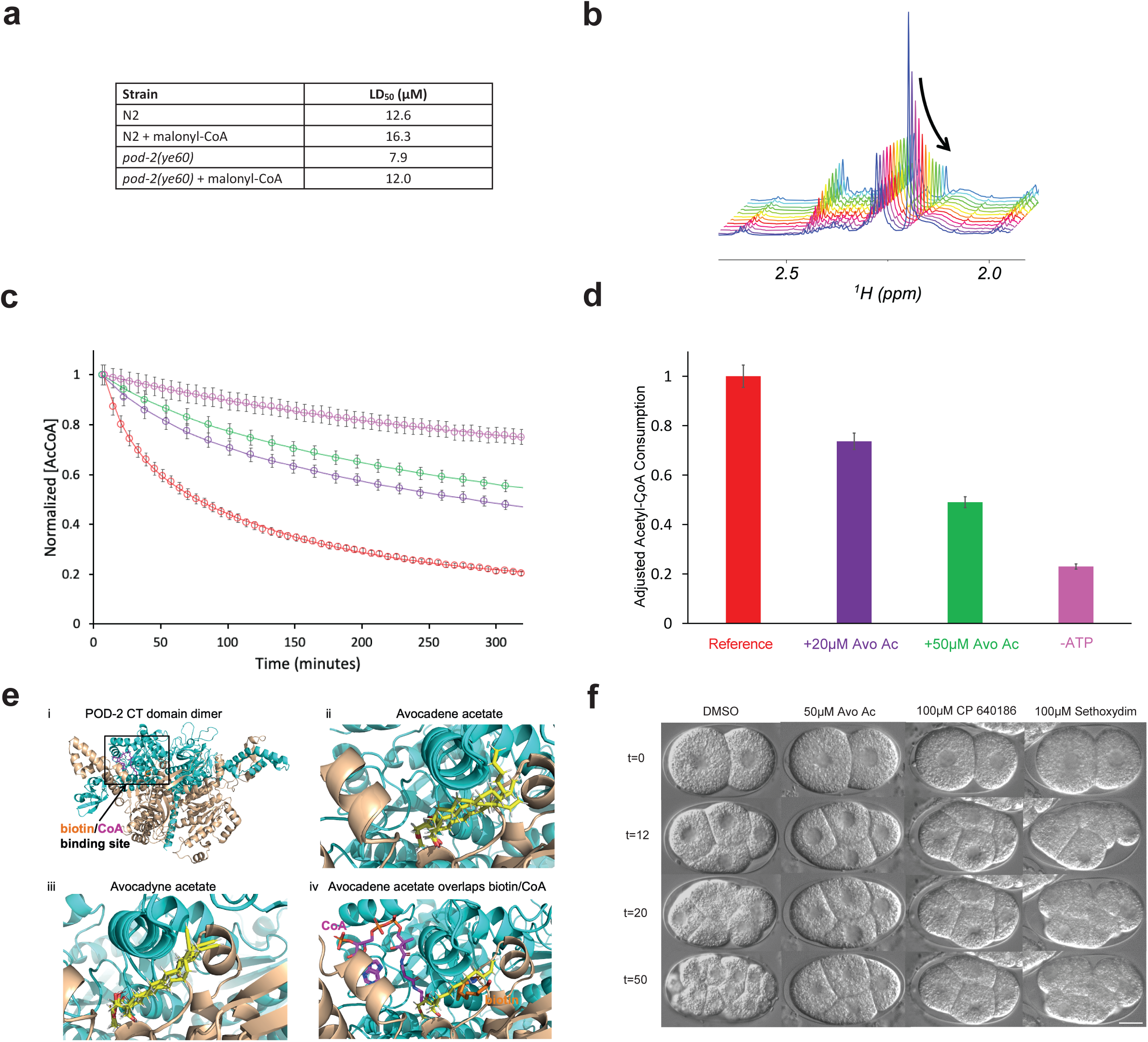
AFAs target fatty lipid metabolism. **a,** LD_50_ values of avocadene acetate in *pod-2(ye60)* and N2 control (L4 stage), in presence or absence of 50 µM malonyl-CoA supplementation. Data represent three biological replicate (*n* = 3) experiments with 4 technical replicates (*n* = 50 animals) each. **b,** ^1^H NMR stacked plot highlighting the decrease of the acetyl methyl peak of acetyl-CoA (AcCoA) at 2.2 ppm, after addition of a *C. elegans* protein fraction containing POD-2 to a standard reaction mixture (“Reference”: 50 µM AcCoA in D_2_O + 50 µM ATP + 30 mM (NH_4_)HCO_3_). **c**, Time course of the *t*_0_-normalized AcCoA concentrations assessed from NMR spectra after POD-2 addition. The reported curves refer to the normalized AcCoA concentration under 4 different composition conditions, namely Reference solution (red), the Reference with 20 µM (purple) or 50 µM (green) avocadene acetate (Avo Ac), and the Reference devoid of ATP (violet). Each curve reports the signal intensity data from a single sample (*n* = 1). The error bars represent the maximal experimental uncertainty affecting the NMR signal areas. **d**, NMR-based consumption of AcCoA relative to the Reference solution, measured 180 minutes after POD-2 addition. The color code and the error bars are the same as in panel c. Each bar represents a single sample (*n* = 1) with the complement to unity of the 180-minutes concentration data-point reported in panel c, after normalization over the corresponding value of the Reference solution. **e,** Structural modeling of avocadene acetate and avocadyne acetate complexed with POD-2. (i) Model homodimer of POD-2 CT domain (residues 1438-2165) with biotin and CoA in the active site (boxed). Dimer and ligand bind sites were modeled using solved yeast CT dimer complexes (PDB ID: 1w2x, 5csl). (ii, iii) Top 3 conformations, as ranked by affinity, of (ii) avocadene acetate and (iii) avocadyne acetate complexed with POD-2. (iv) Overlap of biotin and CoA with the top avocadene acetate conformation at the CT active site illustrates how avocado-derived compounds block POD-2 activity. **f,** DIC images showing that ACC inhibitors CP-640186 and sethoxydim impair embryonic development (*n* = 10), (scale bar = 10 µm). Source data are provided as a Source Data file.

The electron transport chain is the main source of ROS in the cell, and defects in mitochondrial respiration are known to lead to oxidative stress^28^. We measured ROS levels in L4 stage animals in the presence of avocadene acetate using two different fluorescent reporters: a general ROS sensor, chloromethyl-2’,7’ dichlorodihydrofluorescein diacetate (CM-H2DCFDA), and MitoSOX Red, which specifically senses superoxide within mitochondria. MitoSOX is targeted to the mitochondria-rich pharyngeal bulb upon ingestion by *C. elegans,* where it displays red fluorescence upon oxidation by superoxide^29^. In comparison with DMSO-treated controls, the CM-H2DCFDA sensor showed a 30% increase in ROS (unpaired t-test, *p* < 0.0001) (Fig. 4e), while MitoSOX showed a 50% increase in fluorescence intensity in the posterior pharyngeal bulb region (unpaired t-test, *p* = 0.0125) (Fig. 4h,i). The elevated levels of ROS observed using both assays indicate that treatment with avocadene acetate leads to either increased production or impaired neutralization of ROS.

We next explored whether oxidative stress is an important contributor to the anthelmintic effect of AFAs by pre-treating animals with antioxidants. Since glutathione (GSH) is a well-known antioxidant and a substrate of several antioxidant enzymes, we pre-treated worms with the GSH precursor N-acetylcysteine (NAC), which both promotes production of GSH and neutralizes certain oxidizing species directly^30^. L4 worms pre-treated with 10 mM NAC showed a 40% reduction in avocadene acetate-induced larval lethality compared to controls (one way-ANOVA, p-value < 0.0001) (Fig. 4j, Supplementary Fig. 6a,b). We also pre-treated animals with the essential vitamin B12, which has been shown to reduce ROS levels in both the cytoplasm and mitochondria^31^. Vitamin B12 deficiency leads to oxidative stress^31^, and supplementation with vitamin B12 improves mitochondrial homeostasis in worms^32^. We found that pre-treatment with vitamin B12 alleviates the effect of avocadene acetate-induced lethality (Supplementary Fig. 6c,d). These results support the idea that administration of AFAs leads to oxidative stress and that this contributes to their anthelmintic activity.

Since oxidative stress is associated with mitochondrial damage, we tested whether avocadene acetate causes defects in mitochondrial morphology that could be a sign of mitochondrial dysfunction. Using a *C. elegans* strain that expresses GFP in the mitochondrial matrix in body wall muscle^33^, we found that avocadene acetate-treated L4 stage worms displayed a spectrum of mitochondrial defects (Fig. 4d,e). Whereas mitochondria normally form an elongated, tubular network, those in treated animals became fragmented and formed isolated spherical structures of varying sizes that appeared to undergo progressive loss, becoming barely detectable in the most extreme cases. The severity of these effects was significantly higher overall in comparison with DMSO-treated controls (Chi-square test *p* = 0.005) (Fig. 4d,e).

### AFAs target ACC/POD-2

The structures of AFA compounds, coupled with their mitochondrial phenotypes and effects on oxygen consumption and ROS, suggested the possibility that they may interfere with enzymes involved in lipid metabolism. Moreover, previous work using acute myeloid leukemia (AML) cells suggested an involvement of avocatin B in lipid metabolism^34–36^ and a direct interaction of avocadyne with the human very long chain acyl-CoA dehydrogenase (VLCAD)^37^. To test this idea in worms, we performed a chemical genetic screen with candidate genes in the fatty acid synthesis (FAS) and fatty acid oxidation (FAO) pathways.

The first set of genes we tested related to the shuttling of fatty acids into the mitochondria. Medium- and long-chain fatty acids (up to C_20_) are first activated by acyl-CoA synthetase in the cytosol (ACS-2 in *C. elegans*) and are then actively translocated into mitochondria by the carnitine shuttle. This system comprises the outer mitochondrial membrane protein CPT1 (carnitine palmitoyl transferase I, which has five homologs in *C. elegans*) and two inner membrane components, carnitine-acylcarnitine translocase (CACT, DIF-1 in *C. elegans*) and carnitine palmitoyl transferase II (CPT2, *C. elegans* CPT-2). Once inside mitochondria, fatty acids undergo multiple rounds of beta-oxidation, producing electron carriers for oxidative phosphorylation and acetyl-CoA, which can be converted to other metabolites and used to extract additional energy via the tricarboxylic acid (TCA) cycle^38–40^.

If AFA compounds interfere with fatty acid metabolism, then disrupting the shuttle pathway would impact AFA-induced phenotypes. Consistent with this expectation, we found that RNAi knockdown of key carnitine shuttle genes – including *acs-2*, the CPT1 homolog *cpt-4*, *dif-1*, and *cpt-2* – partially rescued the effect of 5 µM avocadyne acetate (Mann-Whitney U test, *p* = 0.0271, *p* = 0.0377, *p* = 0.0151, *p* = 0.0039) (Supplementary Fig. 7). However, RNAi of the CPT1 homolog *cpt-1* had no significant effect (Supplementary Fig. 7, Supplementary table 2). These results suggest that lowering the activity of the carnitine shuttle by RNAi can reduce, but not eliminate, sensitivity to AFA compounds.

We next explored chemical genetic interactions of AFAs with a subset of genes involved in downstream FAO reactions. The first step in each cycle of beta-oxidation is mediated by long-, medium-, and short-chain acyl-CoA dehydrogenases. RNAi of 12 known *C. elegans acdh* genes (*acdh-1* to *4* and *acdh-6* to *13*) had no impact on avocadyne acetate lethality (Supplementary table 2), suggesting that if AFA compounds directly inhibit one or more of these, this can be compensated by some level of functional redundancy. We also found no genetic interaction using the *acdh-12(gk5631) C. elegans* mutant, the homolog with highest sequence similarity to the human VLCAD^41^. Similarly, we could only uncover inconsistent effects when testing the propionate pathways, which we could partly explain by the level of B12 in the food (Supplementary table 2). Consistent with the possibility that AFAs are not metabolized, NMR studies showed that C^13^-labeled avocadene acetate was not degraded in *C. elegans* (Supplementary Fig. 8).

In the process of testing genetic interactions with enzymes involved in fatty acid metabolism, we found that the *C. elegans* Acetyl CoA Carboxylase (ACC) *pod-2(ye60)* temperature-sensitive (*ts*) mutant is significantly more susceptible to avocadene acetate treatment than wild type (Fig. 5a, Supplementary Fig. 9a,b), showing an LD_50_ of 7.91 µM vs. 12.57 µM for WT (unpaired t-test, *p* = 0.003). A strain carrying an auxin-induced degron (AID) fused to POD-2 protein (*pod-2*::AID)^42^ was also significantly more sensitive to avocadene acetate: young adults incubated with auxin for 24h to deplete the POD-2 protein showed an LD_50_ of 2.12 µM vs. 15.14 µM in untreated controls (unpaired t-test, *p* < 0.0001) (Supplementary Fig. 9c). POD-2, like all carboxylases, is covalently biotinylated by BPL-1 (biotin protein ligase 1) to catalyze the ATP-dependent, rate-limiting first step in fatty acid synthesis through the carboxylation of acetyl-CoA to malonyl-CoA^43,44^. Consistent with AFAs acting through inhibition of POD-2, reducing BPL-1 levels by treating *bpl-1*::AID worms with auxin^42^ enhanced the AFA effect (LD_50_ of 8.1 µM vs. 15.05 µM in untreated controls) (unpaired t-test, *p* < 0.05) (Supplementary Fig. 9c).

Malonyl-CoA, the product of the ACC/POD-2 enzyme, plays a major coordinating function in both synthesis and oxidation of fatty acids. If AFAs inhibit the synthesis of malonyl-CoA, exogenous addition of malonyl-CoA could counteract their effect. Indeed, malonyl-CoA supplementation lowered avocadene acetate toxicity in L4 stage animals in both wild type and *pod-2(ye60)* mutants, shifting the LD_50_ from 12.57 µM up to 16.30 µM in WT and from 7.9 µM up to 12 µM in a *pod-2(ye60)* strain (Fig. 5a, Supplementary Fig. 9a,b).

Human ACC, which is conserved by sequence alignment and structural modeling with POD-2 (Supplementary Fig. 10), is reported to be inhibited by the same AFAs *in vitro*^45^. We therefore tested whether AFAs could directly inhibit *C. elegans* POD-2 by purifying a POD-2::6xHis-tagged protein from worms and measuring the enzymatic activity by real-time ^1^H NMR. POD-2 converts acetyl-CoA to malonyl-CoA in an ATP-dependent manner, and the addition of avocadene acetate to the standard (“Reference”) reaction mixture significantly inhibited acetyl-CoA depletion in a concentration-dependent manner (*p* < 0.0001 in any Wilcoxon paired test with Reference alone) (Fig. 5b,c,d, Supplementary Fig. 11 and 12, Supplementary table 3). Because of the keto-enolic equilibrium and the ensuing deuteration in D_2_O, we could not quantify the appearance of the malonyl moiety methylene protons using NMR. However, the production of malonyl-CoA could be qualitatively confirmed by mass spectrometry (Supplementary Fig. 13).

POD-2/ACC functions as a homodimer via a two-step process guided by the biotin carboxylase (BC) and carboxyltransferase (CT) domains to carboxylate acetyl-CoA into malonyl-CoA^46^. Given that the BC and CT domains are both affected by other ACC inhibitors^46^, we performed docking-based *in silico* searches for targets of avocado-derived compounds in both domains (Methods). Using structural alignment with the solved yeast ACC complexes (PDB: 1w96, 5csl), we centered our search in the vicinity of the soraphen A and biotin binding sites in the BC and CT domains, respectively. Computational models of avocadene acetate and avocadyne acetate compounds at the CT site (Fig. 5e) predicted affinities (K_d_ values) of 2.53 μM and 1.66 μM, respectively, two orders of magnitude stronger than predicted values at the BC site (∼400 μM). Persin has a similar predicted binding affinity (3.49 μM), whereas avocadynofuran essentially does not bind these areas of POD-2 (>100,000 μM), consistent with the functional results (Figs. 1 and 2). Together, the *in silico* docking results suggest that the CT active site is the primary target of avocadene/avocadyne acetate and persin inhibitors. Due to the position of the acetate group, this analysis could not reveal mutations that would simultaneously maintain enzymatic activity and significantly reduce avocadene/avocadyne acetate binding. Docking analyses also revealed that CP640186 and sethoxydim, two known ACC inhibitors, could potentially act on POD-2 in the same manner. We found that both compounds strongly inhibited embryonic development, and CP640186 essentially phenocopied AFA treatment (Fig. 5f). However, neither CP640186 nor sethoxydim appears to cross the eggshell barrier: drug-induced developmental phenotypes required breaking the eggshell when mounting embryos under a coverslip for microscopy and were not observed in liquid cultures of embryos, nor in larvae. Together these data support the conclusion that these different pharmacological interventions affect the worms in a remarkably similar way, but they differ in their ability to penetrate the worm eggshell and cuticle.

## Discussion

Due to the limited classes of anthelmintic drugs in use, widespread anthelmintic resistance is particularly severe in the livestock industry worldwide and has also been documented in humans and companion animals^6,11^. Here, we present a novel class of related natural lipid alcohols, AFAs, that show anthelmintic effects *in vitro* and *in vivo*. AFAs are effective in eliciting lethality across all life stages tested across multiple species, including a field-derived *H. contortus* multi-drug resistant strain. When tested *in vivo*, Avocatin A treatment caused a significant parasite load reduction of both adults and eggs. We observed no overt host toxicity in mice treated with Avocatin A, consistent with our data in cell lines, as well as published reports describing a favorable safety profile for Avocatin B (the non-acetylated version of Avocatin A) in mice and preliminary results from phase I clinical trials^47^.

Avocado extracts are a rich source of compounds with diverse bioactivities, antimicrobial and medicinal properties, and safety profiles^48,49^. Isolated avocadene and avocadyne were recently shown to have mosquito larvicidal activity^50^. While purified AFAs have not previously been described to affect nematodes, the polyphenols quercetin and epicathecin from seeds of *Persea americana* are reported to affect larvae of the parasitic nematode *H. contortus* in goats^51^, and extracts from the related species *Persea willdenovii* also show anthelmintic properties^52^. Thus, avocado fruits appear to have evolved a variety of ways to resist infections from insects, nematodes, microbes, and potentially other pests.

Multiple lines of evidence indicate that AFA compounds exert their anthelmintic effects by interfering with lipid homeostasis in *C. elegans* via the lipid biosynthesis enzyme ACC/POD-2: (1) The enzymatic activity of purified POD-2 is directly inhibited by AFAs. (2) Depleting POD-2 or its upstream biotin ligating protein BPL-1 enhances AFA-induced lethality. (3) Because POD-2 converts acetyl-CoA to malonyl-CoA, the rate-limiting substrate for de novo fatty acid biosynthesis, targeting either POD-2 or BPL-1 depletes the endogenous malonyl-CoA pool. Consequently, supplementation with malonyl-CoA in the presence of AFA partially rescues worm development and survival. (4) Malonyl-CoA also serves a key role in balancing lipid biosynthesis and catabolism by negatively regulating CPT1 and thus FAO, and RNAi knockdown of several carnitine shuttle genes partially alleviates AFA-induced toxicity.

Lipid metabolism enzymes have been proposed as “chokepoints” for worm survival and potential targets for anthelmintic drug discovery^53^, and recently an insecticide that acts as an ACC inhibitor (spirotetramat) was shown to be active at high concentration against a plant parasitic nematode *Heterodera schachtii* (50mM) and *C. elegans* (200 µM)^54^.

The embryonic defect caused by direct application of AFAs is more puzzling. While we observe an effect in early embryos, POD-2 protein is not reported to be highly expressed in early *C. elegans* embryos; rather, malonyl-CoA is thought to be supplied maternally by the soma to the germline^55^. However, transcriptome analysis of early embryos detects *pod-2* transcripts^56^, which could be maternally inherited. The rapidly dividing cells in the non-feeding early embryo require a substantial supply of lipid entities for new cell membrane formation and for energy production. AFAs and other pharmacological ACC inhibitors, CP640186 and sethoxydim, all halt embryonic development at any stage including in very early (4-cell) embryos, suggesting either that POD-2 activity is inhibited in those stages, or that another molecular target yet to be identified is present in embryos. The POD-2 paralog T28F3.5 is not reported to be expressed in the early stages of embryogenesis^51^ and is thus unlikely to be a major contributor to early embryonic phenotypes. Focusing on the POD-2 CT active site (Fig. 5e), both avocadene acetate (Fig. 5e-ii) and avocadyne acetate (Fig. 5e-iii) are predicted to bind in a location that overlaps both the biotin and CoA binding sites (especially near its sulfur atom) in solved structures (Fig. 5e-iv). The acetyl group of avocadene/avocadyne acetate binds near CoA, whereas the compound’s tail region extends to the biotin site and beyond. Moreover, all top three favorably docked conformations of these AFAs show significant consistency in the position of the acetyl group; most conformational variations are observed near the tail ends. This analysis suggests that the inhibitory mechanism of avocadene/avocadyne acetate differs somewhat from the synthetic compound CP-640186, which competes with biotin binding but not with CoA^57^.

The anthelmintic effect of AFA across a broad range of free-living and parasitic nematodes and their *in vivo* efficacy at treating infected mice make them great candidates for further development as anthelmintics and point to the conclusion that fatty acid metabolism/homeostasis in nematodes is an interesting avenue for future studies of potential anthelmintic targets.

In summary, the AFA compounds described here have potential to serve as new anthelmintic drugs, either alone or in combination with other anthelmintic compounds. Synergy between compounds with different mechanisms of action could decrease the risk of toxicity and potential off-target effects by reducing the concentration of individual compounds needed to achieve potent anthelmintic activity. This study thus opens new possibilities for the formulation of new drug cocktails that could protect against the development of resistance by targeting multiple pathways essential for the development and survival of parasitic nematodes.

## Methods

The research reported herein complies with all relevant ethical regulations. All mice experiments were performed in compliance with any relevant ethical regulations, specifically following an institutionally reviewed and approved animal use protocol as well as the policies and guidelines of the UK Home Office Project Licence approved by the University of Glasgow Animal Welfare and Ethical Review Board.

### Chemical sources

The library presenting 2320 compounds, including FDA-approved drugs, bioactive compounds and some natural products, was purchased from Microsource Discovery Systems, Inc. Additional quantities of individual AFAs were purchased from Microsource Discovery Systems and Sigma-Aldrich. Indole-3-acetic acid, malonyl-coenzyme A, N-acetyl cysteine, serotonin and vitamin B12 were purchased from Sigma-Aldrich. The ACC inhibitors used in the study are CP-640186 (Cayman Chemicals), spirotetramat (Sigma) and sethoxydim (Dr. Ehrenstorfer). NEOBEE 1053 oil for mice experiments was purchased from Stepan. Avocadane acetate and labeled C^13^ Avocadene acetate were synthesized by ChiroBlock GmbH. For each batch of AFA compounds purchased, purity and activity was determined by NMR and LD50 on N2 controls, respectively. Persenone A and persin were extracted and purified from Colombian Hass avocado according to a previously reported procedure^45^. The purification was performed with a 1260 Infinity series Agilent HPLC system (Santa Clara) coupled to a G4212B photodiode array detector (PDA) and self-collector. The PDA detector was set at 235 nm. A Zorbax C18 column (4,6 x 250 mm x 5 µm, Phenomenex) kept at 25°C was employed. The mobile phase compositions were 0.1% trifluoroacetic acid (TFA) in water (A) and 0.1% TFA in CH3CN (B). Solvents were pumped at 1 ml min^−1^ using the gradient program: 0-6 min, 30% B; 6-12 min, 30%-90% B; 12-28 min, 90% B; 28-35 min, 30% B; followed by 5 minutes re-equilibration. Persenone A and persin were eluted at 18.6 and 19.5 minutes, respectively.

### Worm strains

*P. pacificus* PS312 and *C. elegans* N2 Bristol wild-type strains, SJ4103 (zcls14 [*myo-3::GFP(mit)*], CB1370 (*daf-2(e1370)*), HY520 (*pod-2(ye60)*), JD608 (*avr-14(ad1302); avr-15(ad1051) glc-1(pk54)*), CB3474 (*ben-1(e1880)*)*, RB2119* (*acr-23(ok2804)*), CF1038 (*daf-16(mu86)*), NM1968 (*(slo-1(js379)*), PR1152 (*cha-1(p1152)*), VC731 (*unc-63(ok1075)*), CB193 (*unc-29(e193)*), VC4560 (*acdh-12(gk5631[loxP + myo-2p::GFP::unc-54 3’ UTR + rps-27p::neoR::unc-54 3’ UTR + loxP]) II*) were obtained from the *Caenorhabditis Genetics* Center (CGC). Murat Artan and Mario de Bono kindly provided the PHX1772 (*pod-2(syb1772[pod-2::His10]*), AX8465 (*pod-2(db1461[pod-2::degron::3XFLAG]*)II; ieSi57 II) and AX8273 (*bpl-1(db1372[bpl-1::degron(AID)]) ieSi57[eft-3p::TIR1::mRuby::unc-54 3’UTR+Cbr unc-119(+)] II*) strains. Antony P. Page kindly provided the EG175 (*dyf-17 (ox175)*) strain.

### Chemical screening

Newly hatched *C. elegans* and *P. pacificus* larvae cultivated in 96-well plates were treated with either 1% DMSO (as a control) or library compounds (Supplementary data 2). The primary screen was performed using the NYUAD high-throughput screening automated platform at a concentration of 10 µM in 2 biological replicates (N=2) with 2 technical replicates (n=2) each. Briefly, an overnight culture of OP50 *Escherichia coli* was concentrated 2.5-fold using S-complete medium. 0.5 µl of the chemical was dispensed using the Bravo automated liquid handling platform (Agilent) into each well of the 96-well plate, followed by the addition of 30 µl of bacterial suspension using the MultiFlo FX Multi-Mode Dispenser platform (BioTek). Synchronized L1 stage worms in S-complete were added to 96 wells using the Multiflo Fx device. The plates were incubated at 20°C for 5 days for *C. elegans* and 22.5°C for 7 days for *P. pacificus* in Cytomat automated incubators (Thermo Scientific). These conditions allow animals to develop into the adult stage and give rise to newly hatched progeny. Brightfield images were captured using a Cellinsight CX5 high-throughput microscope (Thermo Scientific) and archived in a custom in-house database. Images were inspected by visual comparison of treated and mock-treated animals to identify compounds that affect larval development or survival, adult fertility, paralysis and/or embryonic lethality in the next generation.

Dose-response experiments for AFAs were carried out in a similar manner at 0, 1.25, 2.5, 5, 10, 25, 50 and 100 µM concentrations. Worms of 4 stages were used: eggs, L1 stage worms, young adults and dauers. Eggs were collected by straining a synchronized egg laying population using a 40-micron strainer that captured the adults and allowed the eggs to pass through. L1 stage worms were obtained by bleaching a synchronized egg laying population and allowing the eggs to hatch for 24 hours. Adults were harvested by allowing a synchronized L1 stage population to grow up to the young adult stage. To culture dauers, a few hermaphrodites were transferred to an NGM plate seeded with OP50 *E. coli* and incubated at 22°C. Plenty of dauers developed after 10 days. For *daf-2* dauers, a chunk of worms was transferred onto an NGM plate with OP50 *E. coli* and incubated at 25°C for 48 hours to develop into dauers. Images of eggs, young adults and dauers before treatment and 48 hours post-treatment were captured to quantify embryonic and larval lethality. For L1 stage worms, images 5 days post-treatment were captured to quantify the percentage of larvae that developed to later larval and adult stages. These experiments were performed in quadruplicate in 3 independent experiments. The effect of avocadyne acetate and existing anthelmintics on *C. elegans* and *P. pacificus* egg hatching (n > 100) was quantified after 48-hour treatment at 5 µM and 10 µM on bleached eggs.

Anthelmintic-resistant mutant strains JD608, CB3474, RB2119, CF1038, EG175, NM1968, PR1152, VC731 and CB193 were synchronized and L1s were tested for AFA toxicity in a dose dependent manner. Avocadene acetate LD_50_ for the anthelmintic-resistant mutants were obtained in a dose response manner as described above.

### Phenotype scoring

For the primary screen, phenotypes were classified as follows: larval lethal, larval arrest, slow growth, sterile and embryonic lethal. The extent of phenotypic severity was assigned as weak, medium or strong based on an approximate estimate of the percentage of the initial count. For dose-response and egg hatching experiments, worm/egg counts were noted pre- and post-treatment, to quantify % alive, % developed and % hatched. For RNAi experiments, a score between 0 to 5 was given based on the extent of development, with 0 being 100% arrested/lethal and 5 being 100% developed/alive.

### High-content screening in mammalian cells

U2-OS cells (ATCC® HTB-96™) were cultured in ATCC-formulated McCoy’s 5a Medium Modified, supplemented with 10% fetal bovine serum (F7524, Sigma), 100 U/ml penicillin and 100 µg/ml streptomycin (P4333, Sigma), in a humidified incubator with 5% CO_2_ at 37°C. The day before compound addition, U2-OS cells were trypsinized and cell density was determined using a Countess II Automated Cell Counter (Thermo Fisher Scientific). 2 × 10^3^ U2-OS cells were seeded in each well of a 384-well plate in 32 µl complete culture medium. The next day, 8 µl culture medium containing compounds at different concentrations were added to the cell culture to achieve 1 µm to 100 µm final concentrations using a Bravo Automated Liquid Handling Platform. 4 biological replicates in 4 different plates for each concentration were included in the experiment, with DMSO as control. The plates were briefly centrifuged for 30 seconds at 500 x *g* and incubated for 24 hours before staining. For staining, 20 µl Mitotracker Deep Red (1:1500 dilution, M22426 Thermo Scientific) was added to the cell culture using a Matrix® WellMate liquid dispenser (Thermo Scientific). After a brief centrifuge, the cells were incubated at 37°C for 30 minutes. Next, 20 µl formaldehyde (13% in PBS) was added to the cells for fixation at room temperature for 15 minutes. The fixative was removed, and the fixed cells were washed twice in 60 µl PBS, using a BioStack Microplate Stacker and MultiFlo FX Multi-Mode Dispenser. After washing, cells in each well were treated with 25 µl of staining solution containing Syto14 Green nucleic acid stain (1:2000 dilution, S7576 Thermo Scientific), Wheat Germ Agglutinin_Alexa Fluor™ 555 Conjugate (1:200 dilution, W32464 Thermo Scientific) and Hoechst 33342 (1:4000 dilution, 62249 Thermo Scientific) for 45 minutes at room temperature. After staining, cells were washed twice with 60 μl PBS and wells refilled with 50 μl PBS using the BioTek platform. Plates were sealed with an aluminum foil cover and stored at 4°C until imaging. Stained cells were scanned with the Cellinsight CX7 High-Content Screening (HCS) Platform (Thermo Scientific), with a 20× objective. Compartmental Analysis Bio Application was applied for the image analysis. The nuclei with Hoechst-staining were identified as primary objects (Circ), and simulated cytoplasm (Ring) was created according to nuclear shape and neighboring cells.

### NMR Experiments

NMR spectra were always recorded at 20°C, on the 14.0 T Bruker Ascend instrument of New York University Abu Dhabi Core Technology Platform, equipped with a single-axis gradient cryoprobe operating at 600.19 MHz (1H). The spectra were recorded at 20°C in deuterated chloroform. *C. elegans* embryos (50,000) or L4 larvae (10,000) were treated with 20 µM and 10 µM avocadene acetate, respectively (Sigma). The treatment times were 0.5 and 2 hours for embryos and 0.5, 2, 4 and 8 hours for L4 larvae. Subsequently, the treated and the corresponding control batches were centrifuged to collect the pellets after removal of the supernatant. The pellets were washed 3 times with distilled water prior to extraction. To check for external avocadene acetate complete removal, the intermediate washing was also carried out with 10% aqueous ethanol, leading to the same results as obtained with the triple water washing.

The metabolite extraction from the embryo and larvae pellets was carried out using a variant of the Folch method. The pellet samples, resuspended in a minimal amount of water (100-150 µL) and kept on ice, were sonicated in vials floating in water/ice bath with an ultrasound transducer immersed in the sample (20 irradiation/pause cycles (2 s/5 s), frequency 20 kHz, power 25 Watts, applied with a Qsonica Q125-220 instrument). The sonicated pellets were then transferred in glass vials and vortexed for 120 s after a 2 mL addition of a CHCl_3_/CH_3_OH/H_2_O (8:4:3) mixture. A second sonication step was subsequently carried out for 15 minutes, with the sample vials floating in ultrasonic bath (20 kHz, 40 Watts applied with an Elmasonic P instrument) prefilled with ice and water. The samples were then centrifuged (15 minutes, 1,250 x *g* at 4°C) and the distinct organic and aqueous phases were separated and dried under N2 stream. The dried organic phase, re-dissolved in deuterated CDCl_3_ (Sigma) was ready for NMR analysis, whereas the aqueous phase, devoid of methanol after drying, was lyophilized and redissolved in D_2_O (Sigma) for the NMR assessment.

The samples for NMR kinetic data determinations were prepared by adding to 30 mM (NH_4_)HCO_3_ (Sigma) dissolved in D_2_O microliter aliquots of acetyl-CoA (Sigma) and ATP (Sigma) mother solutions, to reach final concentrations of 50 µM for both species. A sample without ATP was also prepared for the corresponding control experiment. For the assessment of AFA compound inhibitory effect, a few microliters of concentrated avocadene acetate preparation (Sigma) in DMSO-d6 (Cambridge Isotope Laboratories) were also added to the mentioned D_2_O solutions, setting avocadene acetate concentration at 20 µM and 50 µM. Careful adjustments of the NMR acquisition conditions were preliminarily carried out on each specific substrate or substrate and inhibitor D_2_O solution, before adding a 15 µl aliquot of POD-2-containing solution from the pool previously obtained by affinity chromatography and subsequent buffer exchange with deuterated Tris (Cambridge Isotope Laboratories). Typically, acquisitions started after a 6-8 minutes dead-time delay for equilibration and field homogeneity stabilization. The experimental data were collected by issuing automatic consecutive acquisitions of one-dimension ^1^H spectra (128 scans/spectrum requiring 6.1 minutes) for 10-12 hours. The optimized acquisition sequence included residual HOD suppression obtained by pairing WATERGATE elements^58^ in the excitation sculpting mode^59^. The experimental data were processed with Topspin 4.0.6 (Bruker) and the acetyl-CoA acetyl methyl peak (2.20 pm) of each spectrum was carefully integrated using the value of the first acquired spectrum in each series as normalization reference.

### Experiments in worm parasites

All parasitic nematode (*H. polygyrus*, *T. circumcincta* and *H. contortus* (ISE-drug sensitive and UGA-drug resistant)) stages and species were tested by placing either 50 freshly prepared eggs or 50 L1 larvae in microwell and testing as described for *C. elegans* and scoring survival after 24-48 hours. For parasitic nematode egg purification, fresh faeces from an infected host were blended in distilled water, passed through a 250 µm sieve and then centrifuged at 1100 x *g* for 5 minutes. The pellet was then resuspended in saturated NaCl and re-centrifuged for 5 minutes at 1100 x *g* and the top layer of supernatant was retained from a 38 µm sieve, washed in distilled water before following the bleach and culture methods described for *C. elegans*. *T. circumcincta* eggs were isolated from fresh fecal samples collected from naturally infected lambs and purified by salt flotation, cleaned with bleach, placed on OP50 NGM plates for 4 days to hatch. Cleaned embryos or hatched larvae were picked into 100 µl M9 with bacteria and no more than 1% DMS0 (much less – 0.1%) either alone or containing one compound. Scoring of larval or embryonic lethality was performed after 24-hour incubation at 22°C. *H. contortus* infected fecal samples collected from Moredun, strain MHC03 (ISE) were purified by salt flotation and bleaching, then placed on OP50 NGM plates. Samples were aliquoted in 100 µl total M9 and incubated with compound or DMS0 (1%). Scoring was performed after 24-hour exposure at 22°C. *H. polygyrus* samples were similarly collected from infected mouse feces. For the egg assays the above purification was carried out and eggs were washed in M9 buffer, exposed directly to compounds in M9 buffer and hatching was assessed over 2 days.

The lifecycle of *Brugia pahangi* was maintained by serial passage through mosquitoes (*Aedes aegypti*) and jirds (*Meriones unguiculatus*), as described elsewhere^60^. Adults and microfilariae (Mfs) were obtained from the peritoneal cavity and cultured in 2ml of RPMI 1640 media with 2% glucose and 100U/ml penicillin, 100µg/ml streptomycin antibiotics at 37°C with 5% CO2. Each well of a 24 well plate contained either 4 adults or 50 Mfs and were supplemented with either 1% DMSO or 1-50µM Avo A. Worms were scored for viability and motility after 24 hours.

### Forward genetics screens

Forward genetic screens were adapted from previous methods^61^. Wild-type L4 stage parental (P_0_) worms were mutagenized in 50 mM ethyl methanesulfonate for 4 hours. Synchronized L1 progeny from either F1 or F2 generations were dispensed onto 96-well plates containing 40 µM lethal dose of avocadene acetate in OP50 resuspended in S-complete. Worms were dispensed at a density of 50 L1s per well. Plates were scanned by eye under the dissecting scope for viable worms. In total, 2.1 million F1 larvae (i.e 4.2 million independently mutagenized haploid genomes) and 2.5 million F2 larvae (i.e 5 million independently mutagenized haploid genomes) were screened for avocadene acetate resistance. Therefore, a total of 9.2 million mutagenized genomes were screened.

### Mouse Experiments

Animal experiments were conducted under a UK Home Office Project Licence (PPL) PP4096415 approved by the University of Glasgow Animal Welfare and Ethical Review Board. Female C57BL/6JOlaHsd from ENVIGO (now part of INOTIV) were delivered at 7 weeks old. After 7 days of acclimatization, experiments are started at 8 weeks with 10 animals per treatment condition. Mice were maintained at 20**°**C - 24**°**C, 45%-65% humidity and following a 12 hour light/dark cycle. The *H. polygyrus* life cycle was maintained and the L3 larvae obtained as previously described^62^. Female mice, aged 7–8 weeks, were infected by oral gavage with 200 *H. polygyrus* L3 in 200 µl water. Oral drug administration: Mice were given Avocadene acetate (Microsource discovery systems) at 2.5 mg in 200 µl SEDDS emulsion (89% water, 0.2% tween80 (Sigma) and 0.8% NEOBEE (Stephan), 10% DMSO) by gavage from the day 9 of infection, and treatment was repeated on days, 11, 13, 15, 17 and 19. Control mice were treated identically with SEDDS. On day 21, worm and egg counts were performed. The egg burdens of individual mice were assessed by collecting 2–3 faecal pellets for each *H. polygyrus*–infected mouse at the specified time intervals. Faeces were weighed before being solubilized in 2 ml PBS followed by the addition of 2 ml saturated sodium chloride solution. Egg counts were then carried out with the use of a McMaster chamber and the average number of eggs/g faeces calculated per sample. At the end point of each study, the intestinal adult worms were also counted to give the total worm burden of each individual mouse; the small intestinal tissues were recovered, and total worm burdens enumerated with the aid of a dissecting microscope. Counts and statistics were plotted using GraphPad Prism .

### Microscopy

To obtain early embryos, 5-10 young adult worms were dissected in a 7 μl droplet of 1% DMSO or the respective chemical treatment on a coverslip. The embryos were left in the droplet for 5-10 minutes to ensure sufficient exposure to the treatment. The coverslip was then mounted on a 2% agarose pad on a glass slide. DIC images were acquired every 2 seconds for the duration of 1 hour at 63x magnification for all the time lapse movies. The embryos were treated with avocadene acetate, CP-640186 and sethoxydim at concentrations of 50 μM, 100 μM and 100 μM respectively. A droplet of M9 containing 1% DMSO was used as a control. All images were acquired using Leica DMi8 microscope and processed using ImageJ.

### Motility assessment and scoring

Assays were set up by dispensing approximately 10 synchronized young adult worms in a volume of 50 µl M9 buffer and 10 µl of OP50 bacterial suspension in each well of a 96-well flat bottom plate. Worms were treated with different concentrations (10 µM, 25 µM, 50 µM and 100 µM) of avocadene acetate or DMSO control. Plates were shaken thoroughly using the Multiflo Fx and images of entire wells captured by the Cellinsight CX5 high-throughput microscope. The protocol was set up to capture two images of each well, taken 500 ms apart from each other. This was repeated at 20 minute intervals for a duration of 480 minutes. Image analysis and fractional mobility score was adapted from a previous study^63^.

### Electropharyngeogram recordings

The Screenchip^TM^ (InVivoBiosystems) was used to record electropharyngeograms (EPG) and quantify pharyngeal pump frequency in worms. Disposable microfluidic cartridges of size SC40 were used for all experiments. Synchronized young adult worms were washed 3 times with M9 buffer and incubated with 10 mM serotonin at 20°C for 30 minutes, followed by treatment with avocadene acetate or DMSO control for approximately 5 minutes. Worms were loaded into the SC40 cartridge using a syringe and allowed to settle for 30-60 seconds. Their EPG recordings were taken for 60 seconds using the NemAquire software (InVivoBiosystems). The entire process was monitored under the Leica M125C inverted microscope to ensure that the worm was correctly positioned between the electrodes and only one worm occupied the channel at a time. Readings of 15 animals were taken for each treatment. Mean pump frequencies were calculated by the NemAnalysis software (InVivoBiosystems).

### Mitochondrial morphology scoring

*C. elegans* strain SJ4103, expressing mitochondrial matrix GFP in body wall muscles, was used to measure the mitochondrial morphology changes associated with Avo Ac treatment. Animals at the L1 stage were treated with 0.5% DMSO or 5µM Avo Ac for 48 hours. The animals were transferred to 2% agarose pads containing 1mM levamisole, covered with a cover slip, and imaged using a Leica TCS SP8 confocal microscope with a 63× objective and a 488nm scanning laser. Scoring was performed according to a published method^33^.

### Oxygen Consumption Rate (OCR) measurements using the Seahorse XFe96 Analyzer

Synchronized L1 nematodes were treated with 0.05% DMSO or 5 µM avocadene acetate at 20°C, until they developed to the L4 stage (42-44 hours). Animals were washed three times with M9, and 20-25 animals in 20 µl M9 were transferred to each well of a 96-well Seahorse cell culture plate containing 180 µl of M9. The drugs FCCP and sodium azide were preloaded to the cartridge injection ports to a final concentration of 10 µM and 40 mM respectively. Baseline respiration was measured initially for 30 minutes followed by injection of FCCP (to measure maximal respiration) and sodium azide to measure non-mitochondrial respiration (Basal OCR was derived by subtracting the non-mitochondrial oxygen consumption induced by sodium azide from the measured baseline respiration levels). Basal respiration, maximal respiration and spare capacities were determined for each well and normalized to the number of worms per well.

### ROS measurement

Intracellular ROS levels were measured using chloromethyl-2,7’ dichlorodihydrofluorescein diacetate (CM-H2DCFDA). Synchronized L1s were treated with DMSO (0.1%) or avocadene acetate (10 µM) for 48 hours at 20°C. Worms were washed to remove OP50 and 10 - 15 animals were incubated with 50 µM CM-H2DCFDA for 4 hours in a 96-well plate. Fluorescence emitted at 535 nm upon excitation at 480 nm was measured using an EnSpire Multimode Plate Reader. Fluorescence levels were normalized to animal numbers by measuring absorbance at 600 nm. Fluorescence peaked after 4 hours, so absorbance values at this time point were compared among the control and treated groups.

### MitoSOX™ Red staining

Animals at the L1 stage were treated with DMSO (0.1%) or avocadene acetate 10 µM for 42 hours. They were then washed off plates and MitoSOX staining was performed using around 100 animals in 500 µl M9 + cholesterol and 50 µl of OP50. The animals were incubated with 1 µl MitoSOX (final concentration 10 µM) in a rocker at 20°C in the dark for 24 hours. They were washed 3 times with M9, allowed to forage in a fresh OP50 plate for 1 hour and imaging of the pharyngeal region was performed using a Leica DMi8 microscope, using a 63× objective with 100 ms exposure using fluorescent filters 510/580.

### NAC and Vitamin B12 treatment

Approximately 10 L1 stage worms per well in 96-well plates were treated with or without 10 mM N-acetyl cysteine (NAC) until they reached the L4 stage. Animals were then treated with 15 µM avocadene acetate or DMSO and survival was scored after 2 days. Similarly, L1 stage worms were treated in the presence or absence of 64 nM vitamin B12, followed by treatment with 6 µM, 8 µM and 10 µM of avocadene acetate upon reaching L4 stage.

### RNAi experiments

All RNAi clones were obtained from the Ahringer library (Source BioScience) and their sequence confirmed. The experiments were carried out on L1 stage worms using the same method as described in the ‘Chemical screening’ section, using overnight RNAi bacterial cultures induced with 1 mM IPTG for 4 hours at 37°C, instead of OP50. For avocadyne acetate and *acs-2/ cpt-1/ cpt-4/ dif-1/ cpt-2/ acdh-1/ acdh-2/ acdh-3/ acdh-4/ acdh-6/ acdh-7/ acdh-8/ acdh-9/ acdh-10/ acdh-11/ acdh-12/ acdh-13* screens, 5 µM avocadyne acetate was used.

### Malonyl-CoA rescue experiments on WT and *pod-2(ye60)*

N2 and *pod-2(ye60)* cold-sensitive animals were grown at 24°C. When they reached the L3 stage, they were transferred to 15°C for a duration of 16 hours. Subsequently, the animals were exposed to different concentrations of avocadene acetate. Additionally, one experimental set received a supplementation of 50 μM Malonyl-CoA. The surviving animals were then counted after a span of 48 hours.

### Auxin-inducible degron (AID) experiments

AX8465, AX8273 and N2 strains were used in this assay. Age synchronized L1 larvae were grown on OP50 at a density of approximately 10 - 15 worms per well in 96-well plates as described above. Upon reaching the L4 larval stage, 1 mM auxin (indole-3-acetic acid) was added to plates containing AX8273 and incubated for 48 hours, followed by the addition of various concentrations of avocadene acetate. For AX8465 strain, treatment of auxin at a concentration of 50 µM took place at the young adult stage and avocadene acetate was added 24 hours later. Live and dead worms were quantified after 48 hours.

### POD-2 biochemistry

*C. elegans* strain, PHX1772 *pod-2(syb1772[pod-2::His10]) II* was used for the pull down experiments. Worm protein extract was prepared from 500,000 L4 stage worms, grown in liquid culture. The worms were pelleted, washed 3 times with M9 and resuspended in the lysis buffer. The worm suspension was then added dropwise to a pre-cooled mortar with liquid nitrogen. The frozen beads were ground to a fine powder. This powder was thawed again and homogenized using a QSonica sonicator. The sample was then centrifuged at 12,298 x *g* using a Beckman Coulter optima XPN-90 Ultracentrifuge using Type 90 Ti rotor at 4°C. The clear supernatant was then transferred to equilibrated Ni-NTA agarose resin slurry column and the mix was incubated for 1 hour at 4°C. After collecting the flow through, the column was washed 5 times with the wash buffer and the elution of the protein was done using the elution buffer containing 250 mM imidazole. The eluted fraction was then concentrated by passing through a centrifugal filtration column (100 K molecular weight cut off) and the purified fraction was used for NMR assays.

### Mass spectrometry and Proteomics

*Preparation of in-solution digests*. Bradford assay was performed to measure the protein concentration. Thirty µg protein per sample were reduced with 20 mM DTT (dithiothreitol, Sigma), at room temperature (RT), for 1 h and S-alkylated with 50 mM IAA (iodo-acetamide, Sigma), for 1 h in dark room. The remaining IAA was quenched with 20 mM DTT for 1 h in the dark. Prior to digestion, urea (Sigma) was diluted to a concentration of 1 M with 50 mM (NH_4_)HCO_3_ (ABC, Sigma) to prepare the solvent. The proteins were digested with 1:50 (w/w enzyme:protein) MS-grade trypsin/lys-C mix (Thermo Scientific) for 24 h at RT. The digestion was quenched by lowering the pH with formic acid (Fisher) and the sample was desalted with a ZipTip C18 (Merck-Millipore) column. Samples were resuspended in 10 µl of 0.2% formic acid prior LC-MS^2^ (Liquid Chromatography tandem Mass Spectrometry) analysis.

*LC-MS*^2^ *analysis.* The LC was performed on a fully automated EASY-nLC 1200 (Thermo Scientific). The system was fitted with a C18 column (PepMap^TM^ RSLC, Thermo Scientific) with an inner diameter of 75 μm and length of 15 cm. The column was kept at a constant temperature of 40°C. Mobile phases consisted of 0.1% formic acid for solvent A and 0.1% formic acid in 95% acetonitrile for solvent B. Samples were loaded in solvent A. A linear gradient was set as follows: 0% B for 3 minutes and then a gradient up to 35% B in 45 minutes and to 60% B in 60 minutes. A 10-minute wash at 90% B was used to prevent carryover, and a 15-minute equilibration with 0% B completed the gradient. The LC system was coupled to a Q-Exactive high field orbitrap (Thermo Scientific) equipped with an Easy spray ion source and operated in positive ion mode. The spray voltage was set to 1.8 kV, S-lens RF level to 35 and ion transfer tube to 275°C. The full scans were acquired in the orbitrap mass analyzer that covered a mass-over-charge (*m/z*) range of 350–1800 at a resolution of 120,000. The AGC target was set to 3E6 and maximum ion time to 50 ms. The MS^2^ analysis was performed under data-dependent mode to fragment the top 20 most intense precursors using HCD fragmentation. The MS^2^ parameters were set as follows: resolution, 15,000; AGC target, 1E5; minimum AGC target, 8.0E3, intensity threshold, 4.0E5; maximum ion time, 20 ms; isolation width, 1.4 *m/z*; precursor charge state, 2-5; peptide match, preferred; dynamic exclusion, 30 s, and fixed first mass of 80 *m/z*. The normalized collision energy for HCD was set to 32%.

Protein identifications were determined using Thermo Proteome Discoverer 2.4.1.15 (http://thermofischer.com/) software’s with following settings: enzyme, trypsin; instrument, orbitrap; fragmentation, HCD; acquisition, DDA without merging the scans; and precursor and fragment mass tolerance, 10 ppm and 0.2 Da, respectively. Correct precursor was detected using mass only. Peptide identifications were performed using the UniProtKB_SwissSprot database (release 2021_02, https://www.uniprot.org/uniprot) with *C. elegans* taxonomy. The proteome discoverer search tool was used to search for peptides with fixed carbamidomethylation and variable oxidation of methionine (M) and N-terminal acetylation modifications. FDR threshold was set to 1 %, fold change was set to 2, with 2 unique peptides per protein used to filter the identified protein lists.

After undergoing real-time NMR analysis, the samples were lyophilized, resuspended in 50 µl of water and thoroughly mixed at 1:1 volumetric ratio with matrix solution (2,5-Dihydroxybenzoic acid) (Merck KGaA, Darmstadt, Germany). One µl of each sample was deposited onto the MALDI (Matrix Assisted Laser Desorption Ionization) target plate and the spots were left to dry at ambient condition prior FT-ICR-MS (Fourier Transform Ion Cyclotron Resonance Mass Spectrometry) analysis. FT-ICR-MS allows the collection of high-resolution, accurate mass and isotopic fine structure for direct determination of a compound. The measurements were carried out on a Bruker SolariX FT-ICR mass spectrometer (Bruker Daltonics GmbH, Germany). The system consists of an ESI/MALDI dual ion source and is equipped with a 7.0T superconducting magnet (Magnex Scientific Inc., UK). Measurements were carried out in the MALDI positive ion mode. The spectra were acquired with a time domain of 4 mega words over a m*/z* range of 100 to 900. The laser parameters for ion generation were: wavelength, 400 µm; relative power, 55%; shots, 20; focus setting, ‘medium’; frequency, 200 Hz. The plate offset and deflector plate were set at 100 and 220 V, respectively. Data were acquired using ftmsControl software and subsequently processed with DataAnalysis software (Bruker Daltonics). One hundred scans were acquired for each sample.

### Computational modeling

Modeling of POD-2 structures. We used the POD-2 (isoform c) structure predicted by AlphaFold (downloaded from UniProt). Isoform c, with 2165 residues, includes the BC, CT and BCCP domains. The POD-2 dimer and its bound ligands biotin and CoA were modeled by structural alignment using solved yeast ACC homodimer complexes (PDB ID: 1w2x, 5csl). Specifically, the model POD-2 CT domain (residues 1438-2165) deviates from 1w2x and 5csl by 2.8 and 2.5 Å, respectively, suggesting a high degree of structural similarity to solved structures.

We used the Schrodinger software to perform induced-fit docking of chemical compounds to protein receptors^64^. The induced-fit docking method accounts for the flexibility of ligand and protein receptor site conformations to find low binding-energy complexes. The significance of the results is determined by the binding free energy (ΔG) and structural consistency of the low-energy complexes. The experimentally measurable dissociation constant (Kd) can be computed from the binding free energy (ΔG) using the formula Kd = exp(ΔG/RT), where R is the gas constant and T is the temperature (25°C). For avocado-derived compounds, we set the ligand size parameter to 25 Å to ensure that the search space around the target is sufficiently large. Besides this, dockings were performed using default parameters. Molecular structures of the ligands of interest were downloaded from PubChem and then prepared for docking using Schrodinger’s LigPrep module to generate energy-minimized structures. We used the method to reproduce 3 solved ligand/ACC complexes for CP-640186 (PDB ID: 1w2x), ND-646 (5kkn) and soraphen A (1w96). The compound conformations were reproduced with high fidelity (complex RMSD ∼0.2 Angstrom), and the predicted favorable affinities are 6, 36 and 139 nM, respectively, for CP-640186, ND-646 and soraphen A.

### Statistics and reproducibility

For dose-response, egg hatching and combinatorial chemical and RNAi screens, *N*=3 independent biological replicates were performed, with >50 individuals per replicate. Data visualizations and statistical comparisons between groups were performed using Graphpad Prism (v8.3.0) or R. Best fit dose-response curves and LD_50_ values were generated using nonlinear regression analysis. Unpaired two-tailed t-tests were used to compare groups and *p-*value<0.05 was considered as statistically significant.

## Data availability

All data are available in the main text, the Supplementary Information and/or in the Source data file. Source data are provided with this paper.

## Acknowledgements

We thank Dr. Robert White and the NYUAD Core Technology Platforms, in particular Nikolas Giakoumidis, Lawrence Torres and Reza Rowshan, for assistance with the NYUAD high-throughput screening facility, Rachid Rezgui for assistance with microscopy and Marc Arnoux and Nizar Drou for assistance with whole genome sequencing of mutagenized worms. We thank Malcolm Kennedy for insights on the *Filariae* testing strategy, Alan Healy for insightful discussions on the chemistry of small molecules, Don Moerman for sharing information and Perihan Bouran and Tiffany Kilfeather for administrative help. This work was supported by the NYU Abu Dhabi Research Institute to the NYUAD Center for Genomics and Systems Biology (ADHPG-CGSB). AP was funded by the Biotechnology and Biological Sciences Research Council of Great Britain (BBSRC) grant No. BB/R00711X/. CC and RMM were funded by a Wellcome Trust Investigator Award (Ref 219530) and the Wellcome Trust core-funded Wellcome Centre for Investigative Parasitology (Ref 104111). Worm strains were provided by the CGC, which is funded by NIH Office of Research Infrastructure Programs (P40 OD010440).

## Author Contributions Statement

All free-living nematode, human cells, NMR, and biochemistry experiments were performed in the laboratory of K.C.G and F.P. The anthelmintic screen was set up by H.Z.F., F.R. and P.G.C. F.R. carried out the free-living nematode dose-response experiments, RNAi, motility and pharyngeal pumping assays and auxin degron mutants’ study. S.G. performed the OCR, ROS, mitochondrial characterization, and POD-2 purification experiments. Y.M., F.R. and H.Z.F. performed EMS genetics screens for resistant mutants. Y.M. performed anthelmintic mutant screens and time-lapse microscopy. S.G. and Y.M. prepared samples for NMR experiments. G.E., Y.H. and G.B. carried out compounds’ extraction, NMR and Mass Spectroscopy experiments and analyzed all related data. Parasitic worm dose-response experiments were carried out in the laboratory of A.P. In vivo mouse experiments were planned by H.F. and A.P. and performed by C.C. in the laboratory of R.M. Bioinformatics structural modeling was performed by H.H.G. Mammalian cell experiments were performed by S.K and X.X. Y.P conducted statistical analysis. N.R. setup screening databases and phenotype scoring interface. G.B. helped with critical analysis for target identification. The project was conceived by H.Z.F., K.C.G. and F.P. The manuscript was written by H.Z.F., K.C.G. and F.P.

