## Supplementary information for "A new class of natural anthelmintics targeting lipid metabolism"

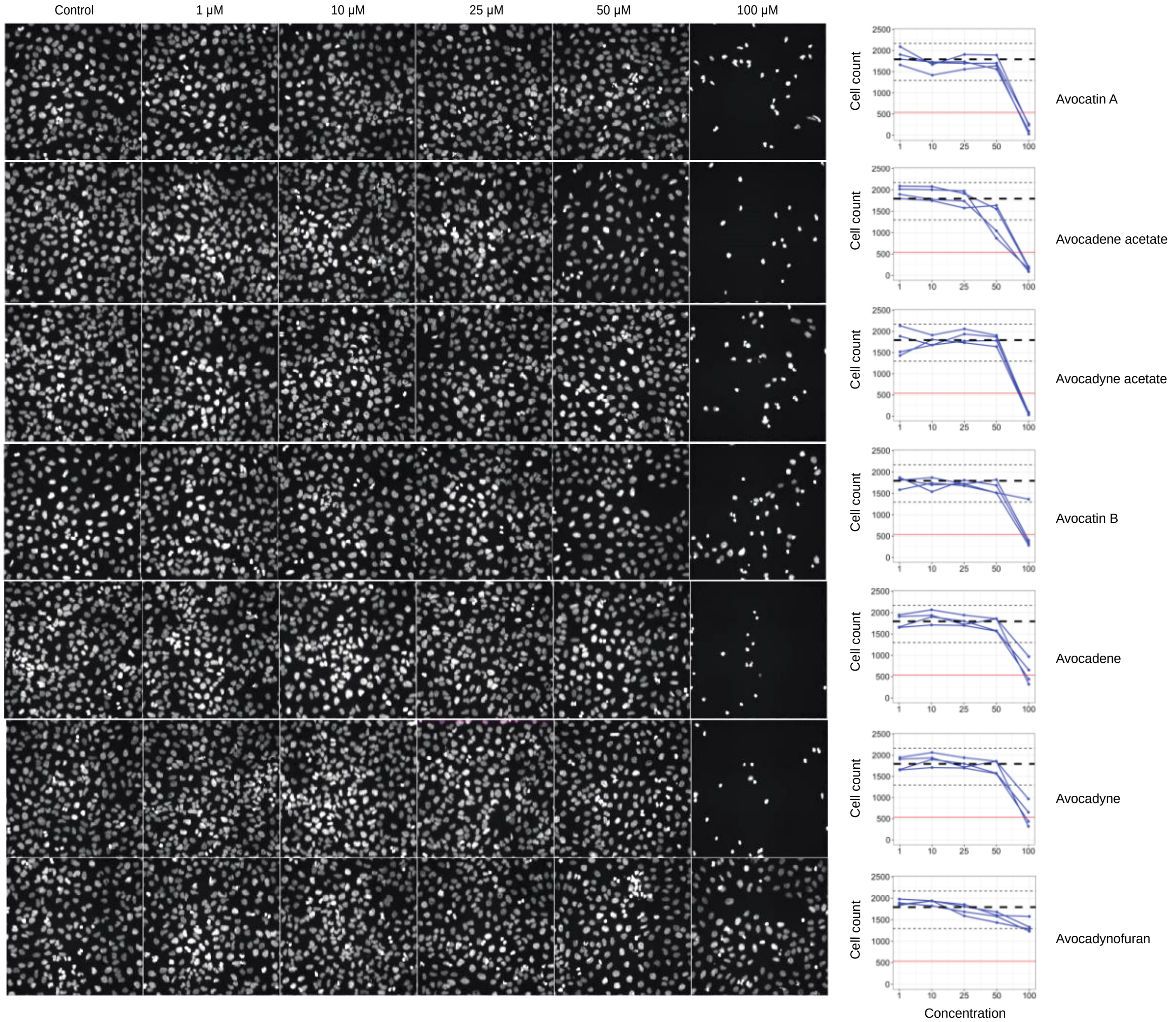


**Supplementary Fig. 1. Dose-dependent responses to AFA compounds in HCS panels.** Left: Representative images of U2OS cells stained with Hoechst to reveal nuclei. Cells were imaged and quantified after 24-hour treatment with control (1% DMSO) or AFA compounds. Right: Dose response curves for replicate assays (blue), showing per-well cell counts (y-axis) vs. compound concentration in µM (x-axis). Median (dashed line) and range (dotted lines, minimum and maximum) of cell counts in DMSO controls and 30% threshold for cytotoxicity (red line) are shown for comparison. Source data are provided as a Source Data file.

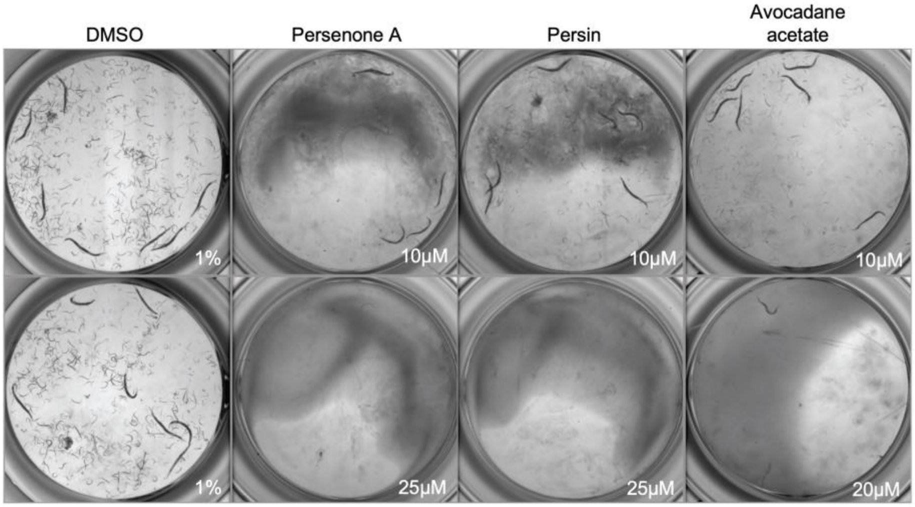


**Supplementary Fig. 2. Persenone A, persin and avocadane acetate impair L1 development of *C. elegans*.** Representative images show the inhibition of development in N2 (WT) L1 stage worms upon treatment with persenone A, persin and avocadane acetate at different concentrations.

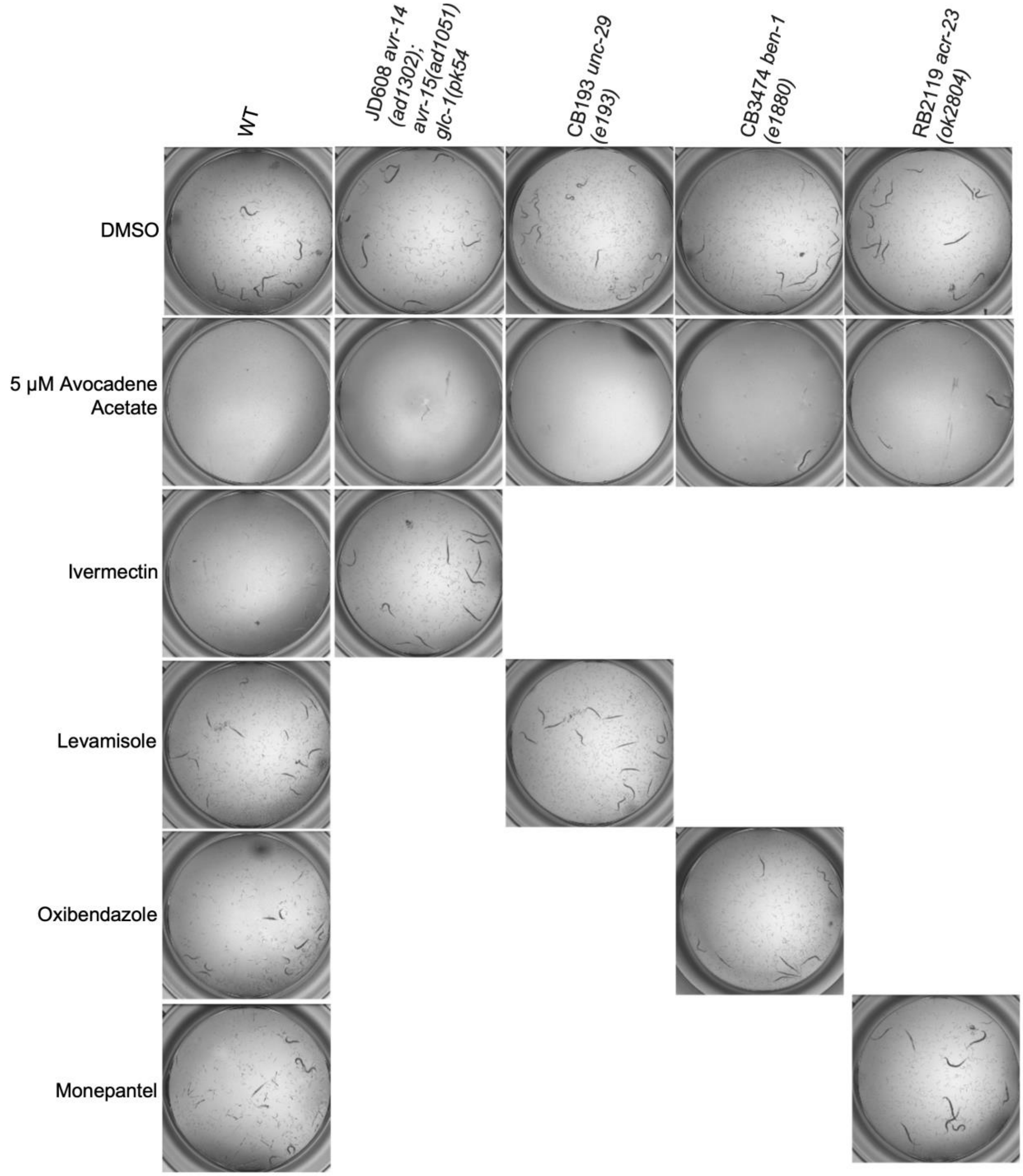


**Supplementary Fig. 3. Anthelmintic-resistant *C. elegans* mutants are sensitive to AFA compounds.** Avocadene acetate is effective against ivermectin-resistant mutant strain JD608 *avr-14(ad1302); avr-15(ad1051) glc-1(pk54),* levamisole-resistant mutant strain CB193 *unc-29(e193)*, monepantel resistant mutant strain RB2119 *acr-23(ok2804)* and benzimidazoles resistant mutant strain CB3474 *ben-1(e1880).*

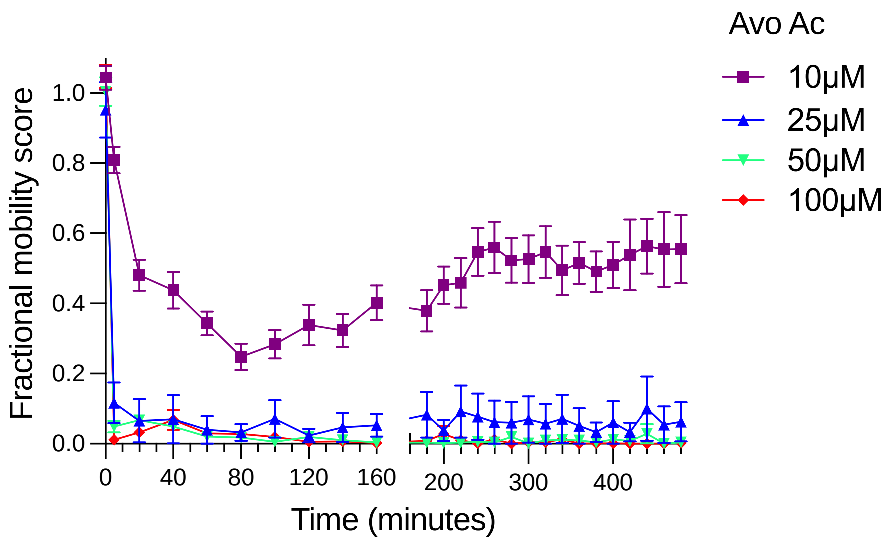


**Supplementary Fig. 4. Kinetic response of young adult *C. elegans* worms treated with varying concentrations of avocadene acetate over a time course of 8 hours.** All data points were normalized by dividing by the fractional mobility score of DMSO control wells for each time point. Data represents mean ± SEM; *n* = 3 BRs; >50 individual worms per concentration. Source data are provided as a Source Data file.

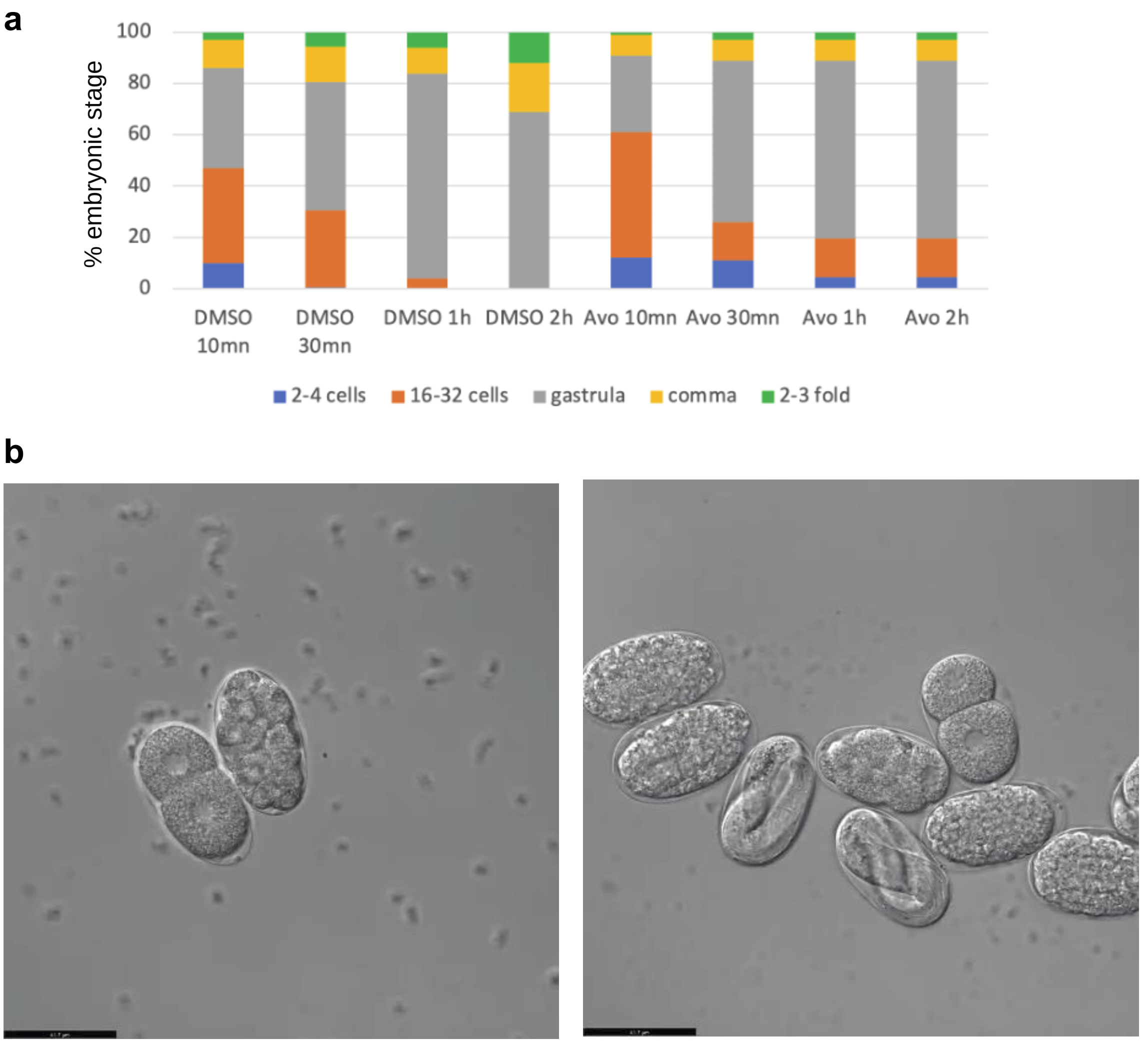


**Supplementary Fig. 5. AFA affects the development of all embryonic stages in *C. elegans*.** (**a**) Stacked bar chart showing relative proportion of different embryonic stages after direct treatment of embryos with DMSO (control) or 20μM avocadene acetate for 10 min, 30 min, 1 hr or 2hr. (**b**) Timelapse videos showing embryo development after treatment with DMSO control (left) or 20 μM avocadene acetate (right). AFA-treated embryos showed developmental arrest within around 20 minutes, *n* = 10, (scale bar = 41.7 μm). Source data are provided as a Source Data file.

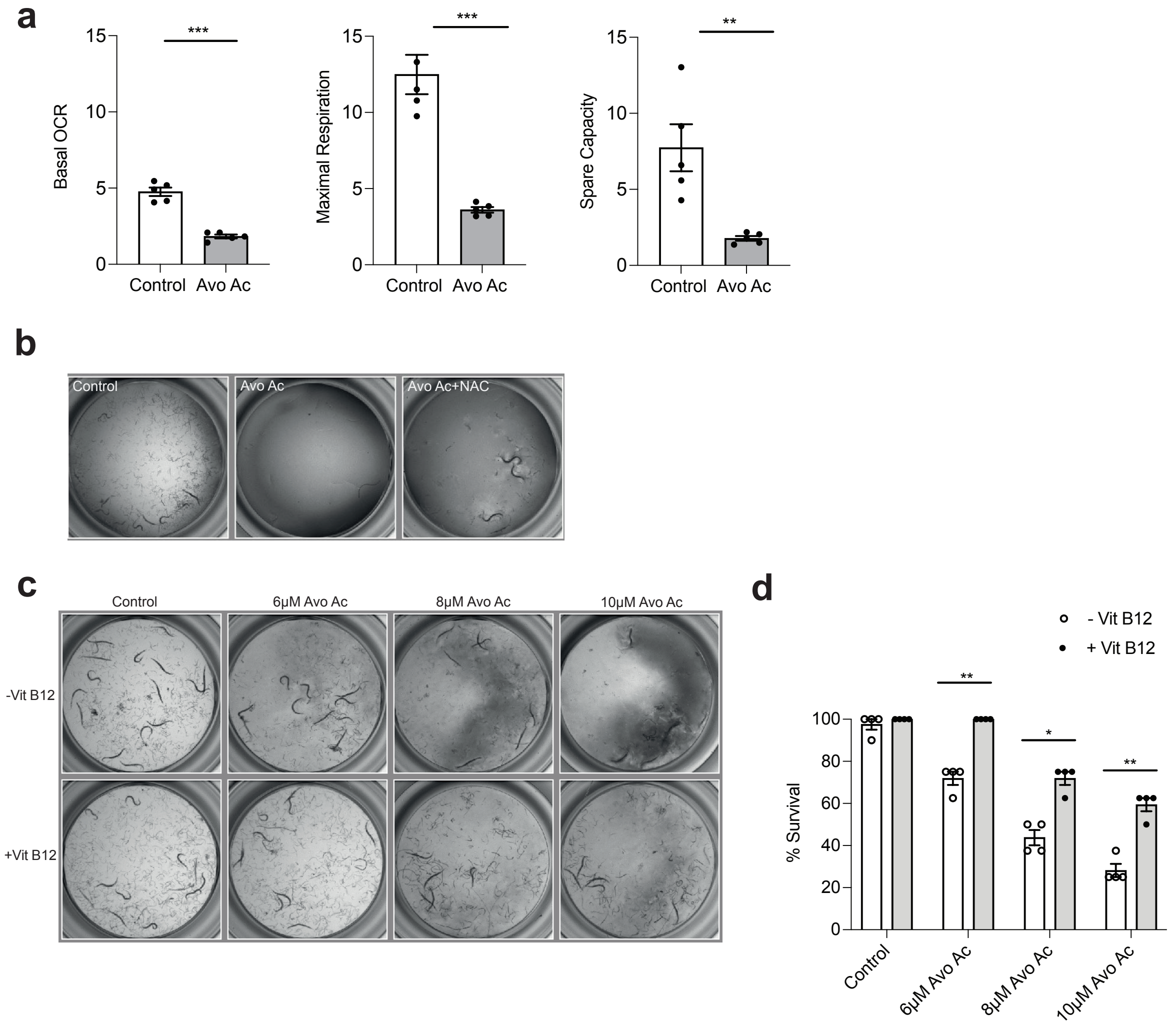


**Supplementary Fig. 6**. **Reduced mitochondrial respiration and lethality induced by avocadyne acetate treatment can be rescued by pre-treatment with NAC and Vitamin B12.** (**a**) Avocadyne acetate treatment impairs mitochondrial respiration. Basal oxygen consumption rate (OCR, left), maximal respiration (middle) and spare capacity (maximal OCR – basal OCR, right) per animal in DMSO controls compared with 5 μM concentration avocadyne acetate treatment of L1 stage animals for 44 hours. $n$ = 3 (**b**) Representative images showing that pre-treatment with 10mM N-acetyl cysteine partially rescued avocadyne acetate-induced lethality. (**c**) Representative images and (d) bar charts illustrating rescue of avocadyne acetate-induced lethality by pre-treatment with vitamin B12. N2 (WT) L1 stage worms were incubated without or with 64nM vitamin B12 until stage L4, followed by treatment with 1% DMSO (control) or varying concentrations of avocadyne acetate for 5 days. Percent development/survival is shown as mean ± S.E.M. ($n$ = 4 BRs; >50 individual worms per assay). Statistical significance (unpaired two-tailed t-test): *$p$-value < 0.05, **$p$-value < 0.001 and ***$p$-value < 0.0001. Source data are provided as a Source Data file.

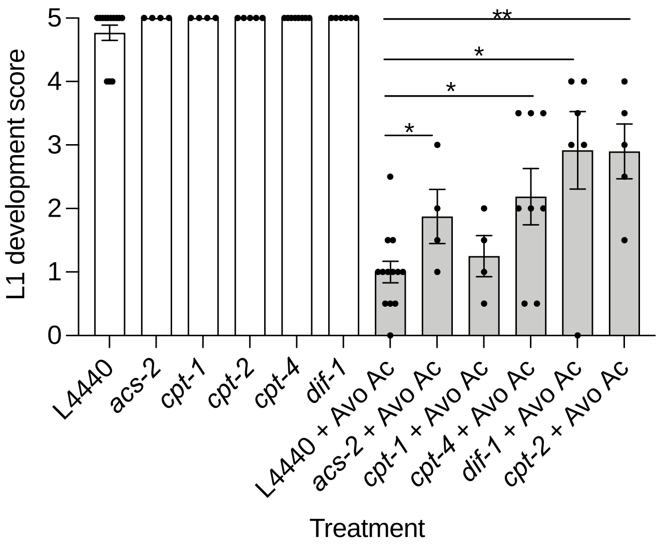


**Supplementary Fig. 7. Knockdown of carnitine shuttle rescues avocadyne acetate phenotype in *C. elegans.*** The bar chart illustrates the effect on development of N2 L1 stage worms grown in the presence of 1% DMSO (empty bars) or 5µM avocadyne acetate (filled bars) and subjected to RNAi by feeding with empty vector (L4440 control) or RNAi of *acs-2, cpt-1, cpt-4, dif-1* or *cpt-2*. Development scores represent the estimated proportion of worms that developed beyond the L1 stage or were alive after 5 days of treatment, with 0 being 100% arrested/lethal and 5 being 100% developed/alive. Shown are mean ± S.E.M.; *n* > 3 BRs; >50 individual worms. Statistical significance (Mann-Whitney U test): *$p$-value < 0.05, **$p$-value < 0.001 and ***$p$-value < 0.0001. Source data are provided as a Source Data file.

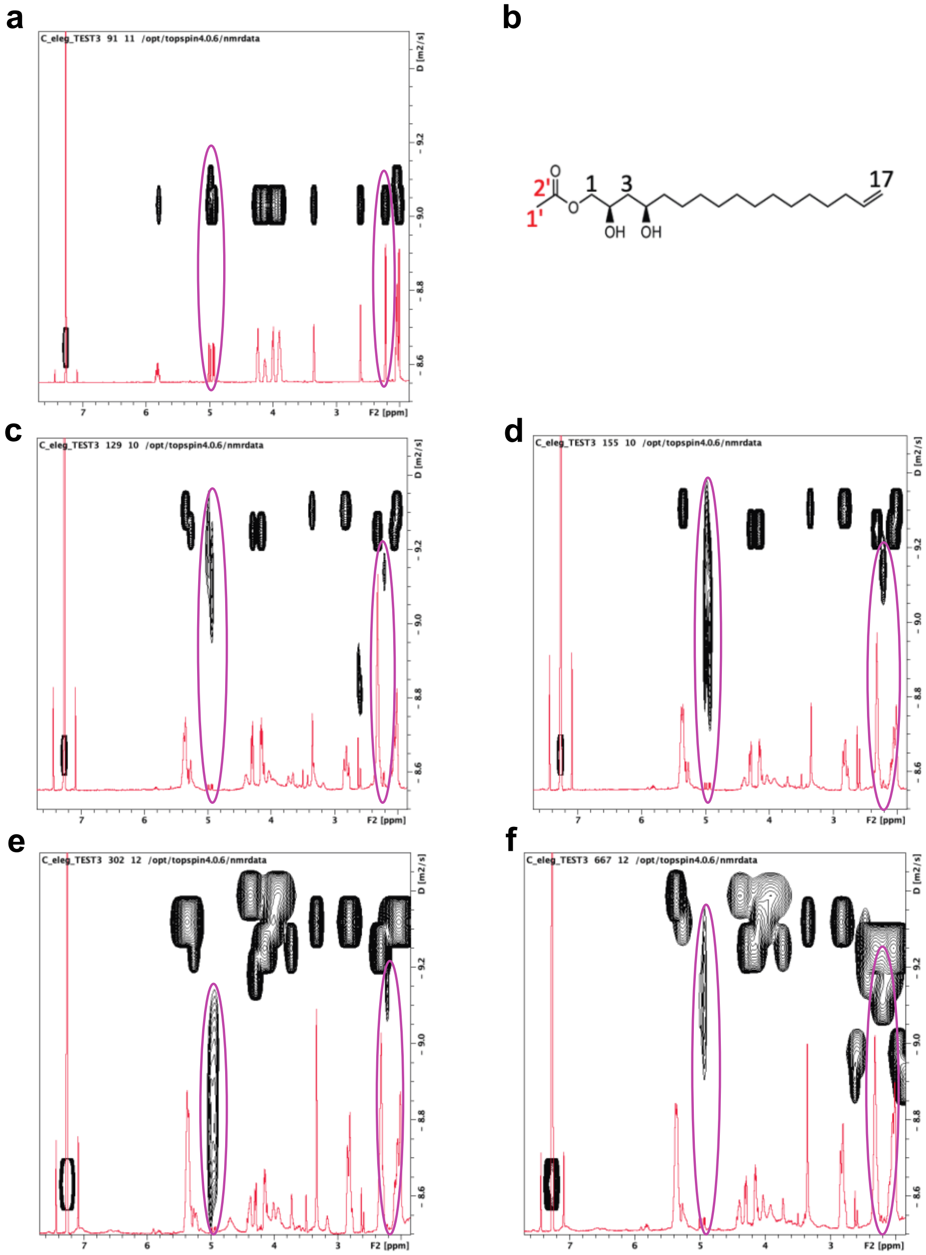


**Supplementary Fig. 8. NMR assessment of molecular integrity of avocadene acetate in whole animal extracts.** (**a**) Molecular structure of avocadene acetate highlighting carbon atoms of the acetyl moiety (1', 2') and the terminal $CH_2$ (17). The molecular species employed for the experiments was synthesized with $^{13}C$ labeling at the acetyl moiety (red labels). (**b-f**) Overlay of $^{1}H$ DOSY map (black contours) and corresponding $^{1}H$ NMR region (red traces) of various samples in deuterium-labeled chloroform ($CDCl_3$) to highlight the uniformity of the translational diffusion coefficient (*D*) of the molecule and rule out hydrolysis giving free avocadene and acetate. Signals from the acetyl methyl (1', ~2.2 ppm) and the terminal $CH_2$ (17, ~5.0 ppm) are circled (purple). (**b**) DOSY map and spectrum overlay for free avocadene acetate in solution. The acetyl methyl (1') and the terminal $CH_2$ (17) exhibit the same *D* value, i.e. $7.66 \pm 0.04 \times 10^{-10}$ $m^2/s$ and $7.56 \pm 0.13 \times 10^{-10}$ $m^2/s$, respectively. (**c-f**) Overlayed DOSY maps and spectra for chloroform extracts of *C.elegans* embryos (**c,d**) or L4 larvae (**e,f**),following (**c,e**) 30 or (**d,f**) 120 minutes treatment with 20 μM or 10 μM avocadene acetate. The measured *D* values for the acetyl methyl (1') and the terminal $CH_2$ (17) of avocadene acetate in the extracts were: (**c**) 6.15±0.50×$10^{-10}$ $m^2/s$ and 5.34±1.92×$10^{-10}$ $m^2/s$, (**d**) 6.26±0.58×$10^{-10}$ $m^2/s$ and 6.43±2.71×$10^{-10}$ $m^2/s$ $m^2/s$, (**e**) 6.23±1.79×$10^{-10}$ $m^2/s$ and 8.56±2.22×$10^{-10}$ $m^2/s$ and (**f**) 5.78±0.41×$10^{-10}$ $m^2/s$ and 5.98±2.81×$10^{-10}$ $m^2/s$, respectively. Despite the errors due to the low intensity of the avocadene acetate signals, the reliability of the determinations is ensured by the observed *D* value of the $CHCl_3$ isotopic impurity of the solvent at ~ 7.2 ppm (invariably 1.90±0.02×$10^{-9}$ $m^2/s$, vs. 2.01×$10^{-9}$ $m^2/s$ expected).

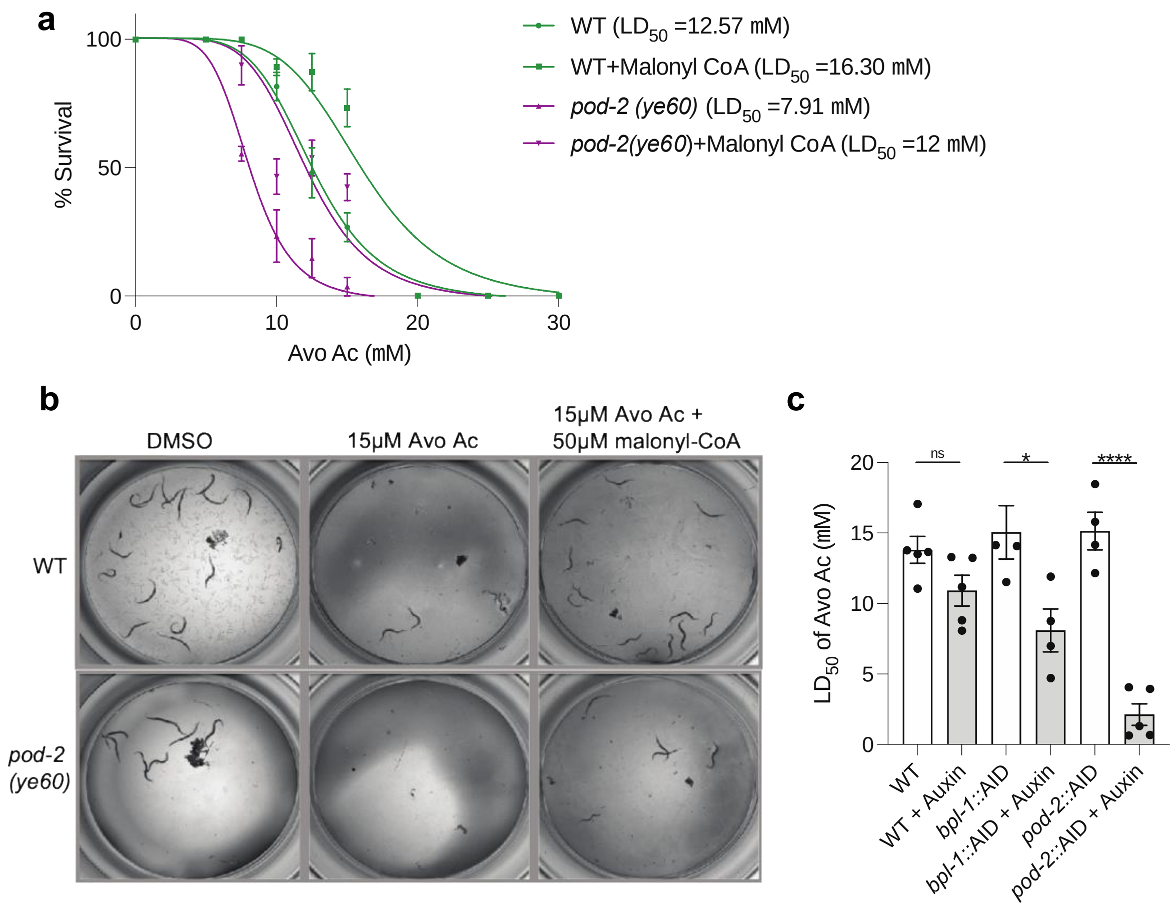


**Supplementary Fig. 9. Malonyl-CoA supplementation alleviates the embryonic lethality phenotype of *pod-2(ye60)* mutant and partially rescues the avocadene acetate lethality phenotype in both N2 and *pod-2* worms**. (**a**) Dose response and LD50 shift curves showing that the *pod-2(ye60)* mutant (purple) is more sensitive to avocadene acetate than WT animals (green) at L4 stage, and that 50 µM malonyl-CoA supplementation partially rescues the lethality. The mean ± SEM of $n$ = 3 BRs with 3 technical replicates per condition are shown. (**b**) Representative images illustrate the partial rescue of the avocadene acetate lethality phenotype by malonyl CoA after 48 hours of treatment, in both WT and *pod-2* mutants. (**c**) Average $LD_{50}$ of wild type, *bpl-1::*AID and *pod-2::*AID strains pre-treated with 1 mM auxin at L4 stage followed by treatment with avocadene acetate at concentrations ranging from 7.5 to 22.5 µM at young adult stage for 48 hours. Each bar in the plot represents mean ± SEM ($n$ > 3 BRs; >50 individual worms per assay). Statistical significance (unpaired two-tailed t-test): ns - not significant; *$p$-value < 0.05; ****$p$-value < 0.0001. Source data are provided as a Source Data file.

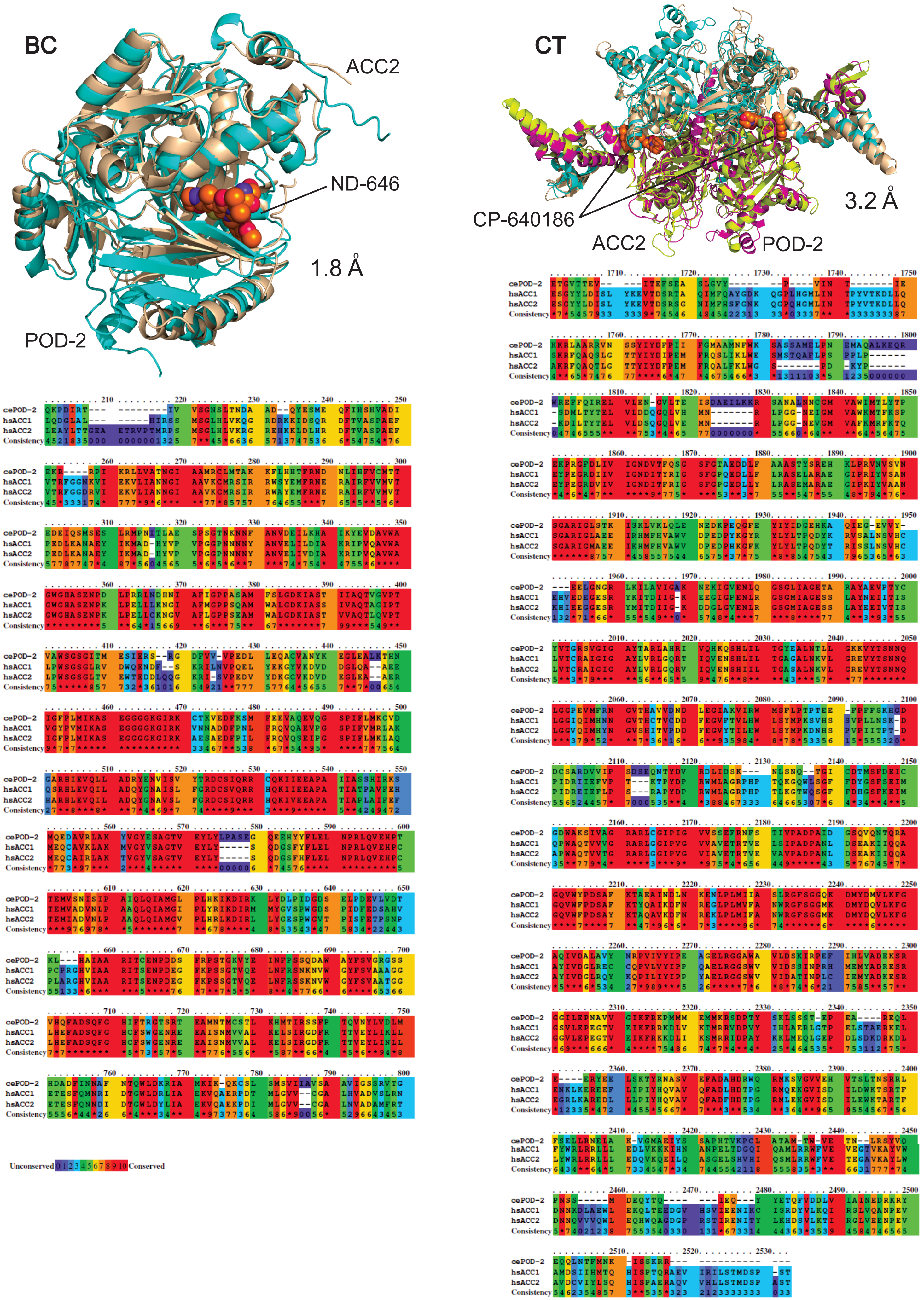
BC
ACC2
ND-646
1.8 Å
POD-2
CT
CP-640186
3.2 Å
ACC2
POD-2
cePOD-2
hsACC1
hsACC2
Consistency
Unconserved 0 1 2 3 4 5 6 7 8 9 10 Conserved

**Supplementary Fig. 10. Comparison of BC and CT domains of *C. elegans* POD-2 and human ACC proteins.** Top: Structural alignments of predicted models for POD-2 BC domain (left) and CT domain (right), superimposed with solved structures of corresponding human ACC2 domains bound to inhibitor compounds. Root mean square deviations (RMSD) between worm and human structures are shown (1.8 Å for BC domain; 3.2 Å for CT domain). The ACC2 BC structure (PDB: 5kkn) contains compound ND-646 at the active site; the ACC2 CT structure (PDB: 3ff6) contains compound CP-640186 at the active site and is shown as a homodimer, with two active sites. Predicted structural models for POD-2 were generated using AlphaFold. Color codes: POD-2, cyan (BC, CT1) and magenta (CT2); ACC2, beige (BC, CT1) and chartreuse (CT2). Bottom: Multiple sequence alignments of corresponding domains from POD-2 with human ACC1 and ACC2.

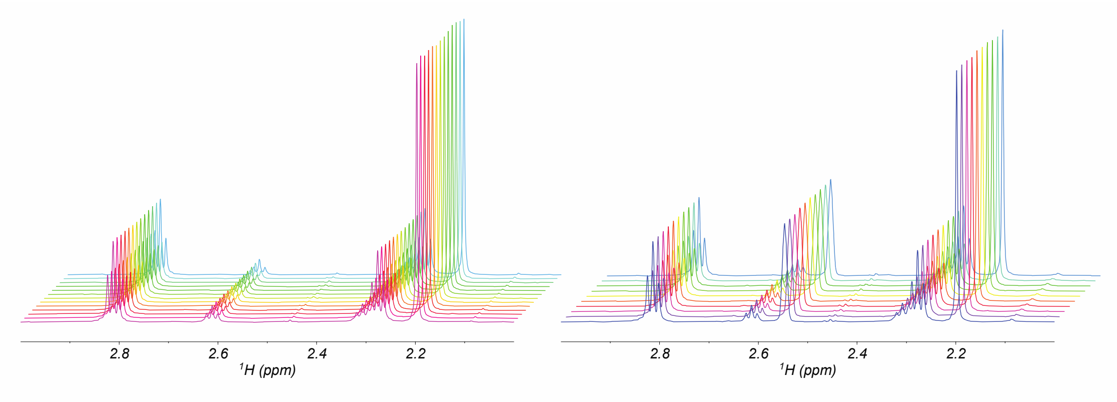


**Supplementary Fig. 11. Stacked plot highlighting the constancy of the acetyl methyl peak of acetyl CoA (AcCoA) at 2.20 ppm in the $^{1}$H NMR spectra** of 50 µM AcCoA dissolved in $D_2O$ with 50 µM ATP and 30 mM $(NH_4)HCO_3$, without (left) and with (right) 50 µM avocadene acetate. The time intervals spanned by the displayed plots are 2.5 hours (left) and 1 hour (right).

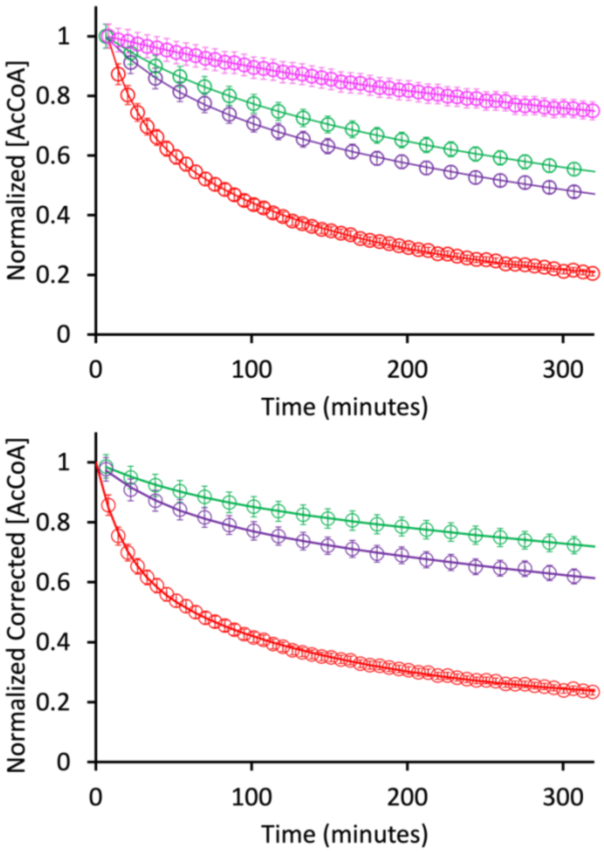


**Supplementary Fig. 12. Time course of the acetyl-CoA normalized concentrations [AcCoA] assessed from NMR measurements after addition of purified POD-2.** The data represent the experimental trends of the standard reaction mixture referred to as reference – 50 µM AcCoA in D2O + 50 µM ATP + 30 mM (NH4)HCO3 – (red), the reference solution with 20 µM (purple) or 50 µM (green) avocadene acetate, and the reference solution devoid of ATP (magenta). In the top plot, data are normalized with respect to starting values. In the lower plot, the three remaining datasets are shown scaled with respect to the data collected without ATP at each timepoint. The difference between any of the reported curves with respect to the reference ones is significant ($p < 0.0001$, Wilcoxon matched-pairs signed rank test). The AcCoA reduction in the absence of ATP is due to mitochondrial AcCoA-acetyltransferase and AcCoA-hydrolase, according to the proteomic assessment of the NMR samples (not shown).The error bars represent the signal-area maximal uncertainty.

The double-exponential fitting curves follow the equation:

$$y = A_0 + A_1 e^{-k_1 t} + A_2 e^{-k_2 t}$$

or, for the solution without ATP, the equation:

$$y = A_0 + A_1 e^{-k_1 t}$$

where $t$ is time, $A_0$, $A_1$ and $A_2$ are pre-exponential constants, $k_1$ and $k_2$ are slow and fast kinetic constants featuring the minimal empirical model fitting the experimental data (see Table S4).

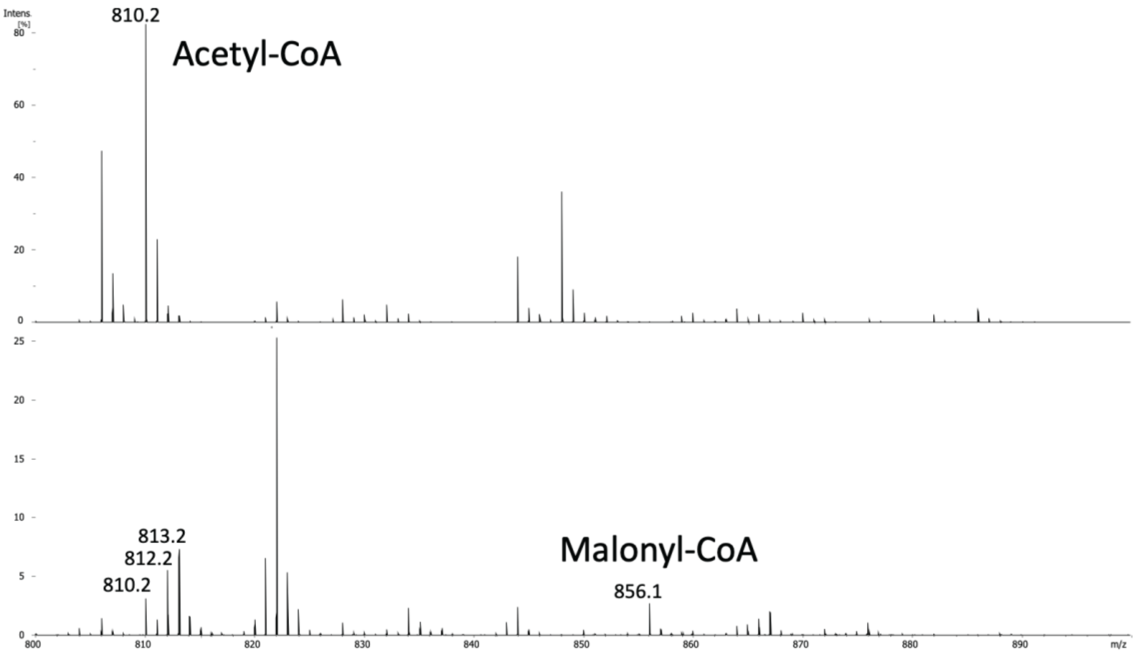


**Supplementary Fig. 13. MALDI-FTICR mass spectra of reference solution after completion of NMR acquisitions** without (upper trace) and with (lower trace) a *C. elegans* protein fraction enriched for POD-2. The reference solution contains the necessary substrates for conversion of acetyl-CoA (AcCoA) to malonyl-CoA (MalCoA) and consists of 50 µM AcCoA, 50 µM ATP, and 30 mM $(NH_4)HCO_3$ in $D_2O$. The upper trace shows the AcCoA $[M+H]^+$ molecular ion at 810.2 Th (Th = thompson, i.e. mass-over-charge ratio unit), as expected without the conversion catalyzed by POD-2. The lower trace contains the weak $[M+H]^+$ molecular peak of MalCoA, at 856.1 Th, as expected from POD-2 activity, along with a peak of unconverted AcCoA at 810.2 Th. Note that the MalCoA molecular ion occurs with an increment of 2 Th because of the incorporation of 2 deuterium atoms from the keto-enol equilibrium in $D_2O$. The low intensity of MalCoA $[M+H]^+$ molecular ion is related to inherently weak ionization and fragmentation, with a step involving $CO_2$ neutral loss leading to AcCoA. The fragmentation-derived AcCoA occurs at 812.1 and 813.1, with the incorporation of 2 or 3 deuterium atoms, which is the signature of the malonyl precursor in 169 D2O.

| *Haemonchus contortus* | | Total eggs | % emb lethality | % larval lethality |
|---|---|---|---|---|
| DMSO | 1% | 857 | 40 | 10 |
| Avocatin A | 10uM | 215 | 77 | 100 |
| | 100uM | 196 | 97 | 100 |
| Avocadene acetate | 10uM | 237 | 78 | 100 |
| | 100uM | 210 | 96 | 100 |
| Avocadyne acetate | 10uM | 207 | 76 | 100 |
| | 100uM | 256 | 81 | 100 |
| Avocatin B | 10uM | 212 | 58 | 78 |
| | 100uM | 238 | 61 | 100 |
| Avocadene | 10uM | 148 | 52 | 75 |
| | 100uM | 204 | 56 | 100 |
| Avocadyne | 10uM | 187 | 39 | 98 |
| | 100uM | 196 | 51 | 100 |

**Supplementary Table 1. AFA compounds impair survival of the parasitic nematode *Haemonchus contortus*.** Embryonic and larval lethality counts after treatment with different AFA compounds in comparison with 1% DMSO control.

| Genes | RNAi + AFA | Mutant + AFA | Mutant genotype |
|---|---|---|---|
| Mitochondrial β-oxidation | | | |
| *acs-2* | rescue* | | |
| *cpt-1* | no effect | | |
| *cpt-4* | rescue* | | |
| *dif-1* | rescue* | | |
| *cpt-2* | rescue** | rescue** | *him-9(e1487) ii; unc-24(e138) let(t...)/nt1 [let(m435)] iv; cpt-2(t1879); dpy-11(e224)/nt1 [let(m435)] v* |
| *acdh-2, acdh-3, acdh-4, acdh-6, acdh-7, acdh-8, acdh-9, acdh-10, acdh-11* | no effect | | |
| *acdh-12* | no effect | no effect | *acdh-12(gk5631)* |
| *acdh-13* | no effect | | |
| *ech-1.1* | no effect | no effect | *ech-1.1(gk5134[loxp + myo-2p::gfp::unc-54 3' utr + rps-27p::neor::unc-54 3' utr + loxp]) v.* |
| *ech-1.2* | no effect | | |
| *ech-2* | no effect | | |
| *ech-4* | no effect | | |
| *ech-5* | no effect | | |
| *ech-7* | no effect | | |
| Propionate/shunt pathway | | | |
| *pcca-1* | variable | | |
| *pccb-1* | variable | | |
| *mce-1* | variable | | |
| *mmcm-1* | variable | enhancement* | *mmcm-1(ok1637) iii.* |
| *acdh-1* | variable | not significant enhancement | *acdh-1(ok1489) i.* |
| *ech-6* | no effect | | |
| *hach-1* | variable | | |
| *alh-1* | variable | | |
| Peroxisomal β-oxidation | | | |
| *acox-1.1* | no effect | | |
| *ech-3* | no effect | | |
| *ech-8* | no effect | | |
| *ech-9* | no effect | | |
| Uncoupling protein | | | |
| *ucp-4* | variable | variable | *ucp-4(ok195) V.* |
| Lipid synthesis and storage | | | |
| *pod-2* | | enhancement** | *pod-2(ye60) ii.* |
| *mlcd-1* | rescue* | | |
| *bpl-1* | | enhancement* | *bpl-1(db1372[bpl-1::degron(aid)]) iesi57[eft-3p::tir1::mruby::unc-54 30utr+cbr unc-119(+)] ii* |
| *fasn-1* | enhancement* | | |
| *emb-8* | | enhancement* | *emb-8(hc69) iii* |
| *c32e8.9 (echdc1)* | no effect | | |
| *fat-5* | no effect | | |
| *fat-6* | no effect | | |
| Other biotin binding carboxylases | | | |
| *mccc-1* | no effect | | |

**Supplementary Table 2. Chemical genetic interactions screen reveals a modified effect of RNAi knockdowns in the presence of AFA.** *mild effect. **strong effect.

| Sample | $A_0$ | $A_1$ | $A_2$ | $k_1$ | $k_2$ | *P-value* |
|---|---|---|---|---|---|---|
| Reference | 0.223±0.015 | 0.551±0.041 | 0.394±0.041 | $7.21 \pm 0.80 \times 10^{-3}$ | $5.66 \pm 1.05 \times 10^{-2}$ | $2 \times 10^{-5}$ |
| Reference + 20μM Avocadene Acetate | 0.297±0.126 | 0.550±0.042 | 0.167±0.042 | $1.64 \pm 0.58 \times 10^{-3}$ | $2.21 \pm 0.57 \times 10^{-2}$ | $3 \times 10^{-3}$ |
| Reference + 50μM Avocadene Acetate | 0.166±0.026 | 0.731±0.067 | 0.107±0.067 | $8.57 \pm 3.98 \times 10^{-4}$ | $1.73 \pm 0.35 \times 10^{-2}$ | $5 \times 10^{-3}$ |
| Reference – ATP | 0.604±0.009 | 0.396±0.144 | – | $3.27 \pm 0.14 \times 10^{-3}$ | – | $1 \times 10^{-3}$ |

**Supplementary Table 3. Fitting parameters for the experimental data reported in Supplementary Fig. 12.** The experimental data fitting was obtained using Prism 10 (GraphPad Software LLC). All parameters are reported with the respective 95% confidence interval (profile likelihood). The reference solution was 50 μM AcCoA dissolved in D2O + 50 μM ATP + 30 mM (NH4)HCO3. The reported parameter values refer to the fitting for normalized AcCoA concentration data, further adjusted by scaling with respect to the corresponding data collected without ATP at each timepoint. This further correction was not applied for the normalized AcCoA concentration data collected in the absence of ATP. The p values represent the probability of the experimental data to fit the exponential models by chance and were computed by comparing the experimental data with their corresponding fitting curves using two-sample two-tailed t-tests.
